# Highly multiplexed imaging of biosensors in live cells

**DOI:** 10.1101/2020.12.11.419655

**Authors:** Jr-Ming Yang, Wei-Yu Chi, Jessica Liang, Pablo Iglesias, Chuan-Hsiang Huang

**Affiliations:** Department of Pathology, Johns Hopkins Medical Institutions, Baltimore, Maryland 21205, USA; Department of Biology, Johns Hopkins University, Baltimore, Maryland 21218, USA; Department of Electrical and Computer Engineering, Whiting School of Engineering, Johns Hopkins University, Baltimore, Maryland 21218, USA

## Abstract

Fluorescent biosensors allow for real-time monitoring of biochemical activities in cells, but their multiplexing capacity is severely limited by the availability of spectral space. We overcome this problem by developing a set of barcoding proteins that are spectrally separable from commonly used FRET (fluorescence resonance energy transfer)-based and single-fluorophore biosensors. Mixed populations of barcoded cells expressing different biosensors can be concurrently imaged and computationally unmixed to achieve highly multiplexed tracking of biochemical activities in live cells.

---

To understand the complex regulatory relationship between multiple signaling, metabolic, and other biochemical events in cells, it is often necessary to study their dynamics under various perturbation conditions. Tracking these activities depends on detecting changes in the physical or chemical properties of the molecules of interest. For example, mass spectrometry or immunoblotting have been used extensively to study various molecular events in cells. While these techniques can detect multiple molecular changes in parallel from a single lysed cell sample, characterizing the temporal evolution requires repeated sampling at different time points followed by laborious processing, making it challenging to follow detailed kinetic responses to large numbers of perturbations. Alternatively, genetically encoded fluorescent biosensors offer a versatile tool to continuously monitor a wide range of biochemical activities in live cells while revealing cell-to-cell variability that is not apparent from ensemble measurements ^1,2^. A major drawback of fluorescent biosensors is their limited multiplexing capability due to the broad emission spectra of fluorescent proteins (FPs) and limited availability of spectral space^3^. Efforts have been directed towards expanding the spectral range by developing far-red/infrared fluorophores or replacing two fluorophore biosensors (e.g. those based on FRET) with single-fluorophore designs ^3-7^. Despite these improvements, in general no more than 6-7 biosensors can be imaged concurrently.

Motivated by the need to monitor the kinetics of multiple intracellular activities, we developed a “biosensor barcoding” method for highly multiplexed tracking of fluorescent biosensors. The key idea is to label cells expressing various biosensors with a set of barcodes, which are combinations of barcoding proteins made of different fluorophores targeted to distinct subcellular localizations. To track multiple events in parallel, mixed populations of barcoded cells, each expressing a different biosensor, are imaged in a single microscopy experiment. Activities from different cells with the same barcode are then pooled together to obtain the average temporal profile of the corresponding biosensor (**Fig. 1**). In theory, N different barcoding proteins can create 2^N^ different barcodes, assuming binary expression (i.e. not expressed or expressed). In reality, the expression of barcoding proteins in individual cells is not all-or-none, thus precluding the use of every possible combination. To ensure robust barcode identification, therefore, we only co-express two barcoding proteins that are 1) of different colors; and 2) targeted to different subcellular locations. Using this scheme, the number of barcodes generated from N fluorophore colors targeted to M subcellular locations is C(N,2) · M · (M-1)= N(N-1)M(M-1)/2, where C(n,r)=n!/(r!(n-r)!) represents the combination formula. **Supplementary Table 1** lists the number of barcodes that can be produced by different numbers of FP colors and subcellular locations.

**Figure 1.**
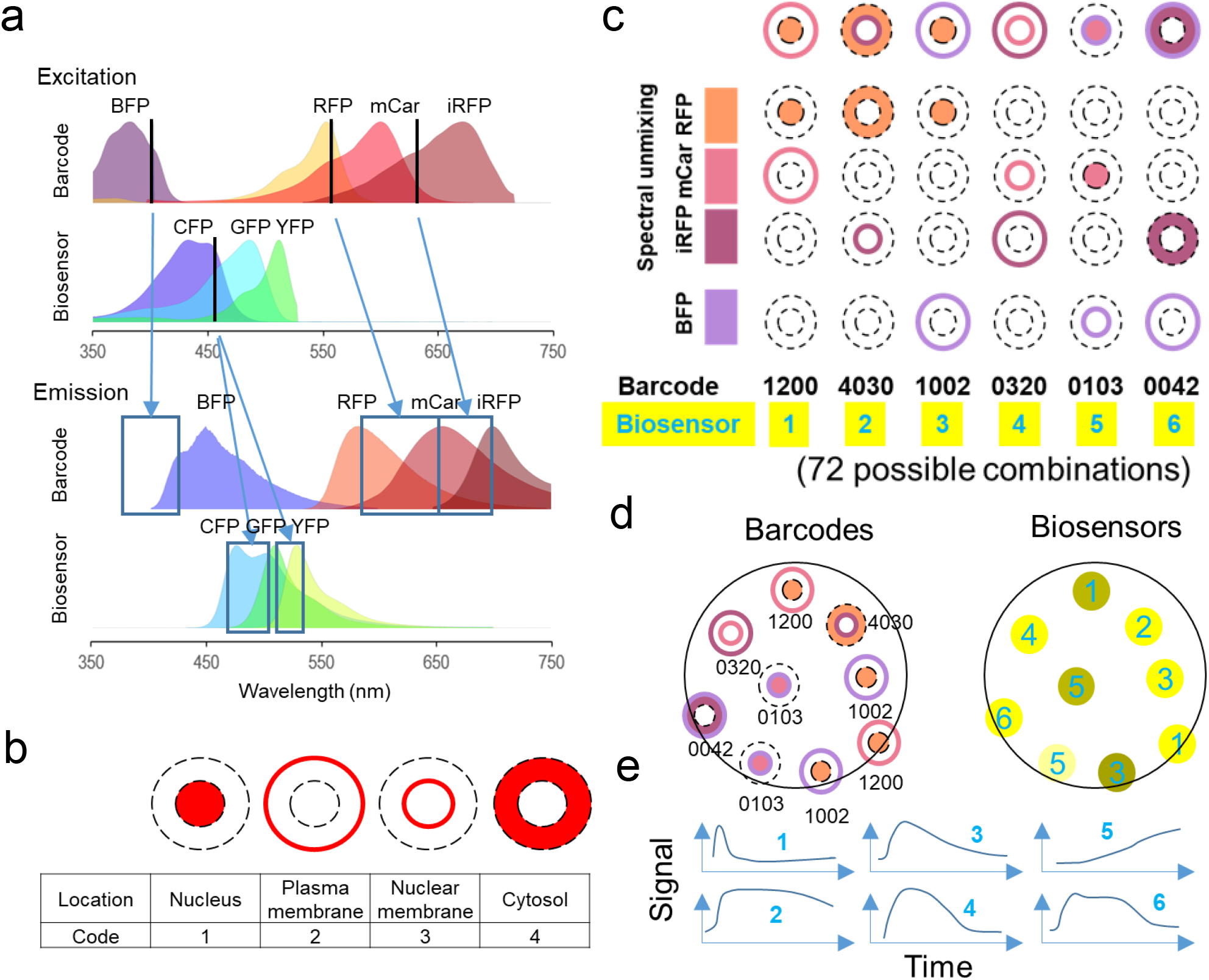
Schematic of Biosensor Barcoding. (a) Excitation and emission spectra of barcoding proteins (EBFP2, TagRFP, mCardinal, iRFP702) and biosensors. Excitation laser lines (405/561/633 nm for barcodes; 458 nm for biosensors) and the corresponding ranges of emission detection are indicated by black lines and boxes, respectively. The profiles of different FPs were generated using the FPbase Spectra Viewer^20^. (b) Targeting sites for barcoding proteins. (c) A pair of barcoding proteins of different colors targeted to two distinct sites are coexpressed with a fluorescent biosensor in the cell. For the sake of simplicity, we use 4 numbers to denote the location of TagRFP, mCardinal, iRFP702, and BFP (in that order), with 1, 2, 3, and 4 denoting the nucleus, plasma membrane, nuclear membrane, and cytoplasm respectively. The number 0 indicates no expression of the FP. For example, the barcode 1200 denotes TagRFP at location 1 (nucleus), mCardinal at location 2 (plasma membrane), and no iRFP or BFP. (d) Cells are mixed and imaged for the barcodes using spectral detectors, and for the biosensors using CFP/YFP channels. The identity of the biosensor can be inferred after spectral unmixing of the barcodes. (e) Activities corresponding to the same barcodes are averaged to obtain the temporal profile of each biosensor.

Our strategy requires that barcoding proteins be spectrally separable from biosensors. A large number of fluorescent biosensors are based on detecting changes in: 1) intracellular localization or intensity of a single fluorophore, in many cases GFP; or 2) the FRET efficiency between a donor and an acceptor, most commonly cyan and yellow FPs (CFP and YFP) ^8,9^. Therefore, the cyan-green-yellow range of the emission spectrum (~450-550 nm) covers a vast number of fluorescent biosensors ^8^. Considering that most commercial fluorescent microscopes detect emissions between 350 and 700 nm, we assessed the effectiveness of FPs in the ranges of 350-450 nm and 550-700 nm as barcoding protein fluorophores.

We first tested whether red/far-red FPs (550-700 nm) targeted to different subcellular sites can be reliably distinguished. To this end, we generated barcoding proteins composed of TagRFP ^10^, mCardinal ^11^, or iRFP702 ^12^ (**Fig. 1a**) fused to localization sequences for: 1) nucleus; 2) plasma membrane; 3) nuclear membrane; and 4) cytoplasm (**Fig. 1b**). To resolve the spectral overlap between fluorophores, we applied the method of linear unmixing of spectral images (see **Methods** for details). Briefly, we took spectral images of cells expressing combinations of these barcoding proteins. Knowing the emission profile of individual fluorophores, the contribution of each can be determined by linear algebra matrix operations. In general, the robustness of linear unmixing reduces with increasing numbers of component fluorophores. We therefore divided the process of unmixing the three fluorophores into two steps: 1) unmixing TagRFP and mCardinal using 561 nm excitation; and 2) unmixing mCardinal and iRFP702 using 633 nm excitation (**Supplementary Fig. 1**). This is feasible due to the minimal excitation of iRFP702 and TagRFP by 561 nm and 633 nm lasers, respectively (**Fig. 1a**). To minimize the discrepancy between the expression levels due to unequal uptake of plasmids, we cloned pairs of barcoding proteins into dual expression vectors (**Supplementary Fig. 2**). As shown in **Supplementary Fig. 3a**, linear unmixing of spectral images of cells transfected with dual expression vectors could be used to robustly resolve the localization and color of the two barcoding proteins.

To further increase the possible number of barcodes, we included BFP (EBFP2) ^13^ as the fourth fluorophore, since: 1) it is not excited by the 458 nm laser used for biosensors; and 2) a properly chosen spectral range (e.g. 400-430 nm) detects the emission from BFP but not CFP in biosensors (**Fig. 1a**). No unmixing was required between BFP and the red/far-red FPs due to their wide spectral separation. Exemplary images of cells co-expressing BFP and one of the three red-far red barcoding proteins are shown in **Supplementary Fig. 3b**. A total of 72 barcodes can be generated using four FPs targeted to four subcellular locations (**Supplementary Table 1** and **Supplementary Fig. 2**).

To test whether biosensor barcoding can achieve highly multiplexed tracking of cellular activities, we mixed barcoded HeLa cells expressing 24 FRET and single-fluorophore biosensors that report the activities of 14 distinct targets in various subcellular compartments, including kinases (AMPK, ERK, p38, JNK, PKC, FAK, Src, PI3K), G-proteins (Gαi1, 2, 3), calcineurin, RhoA GTPase, and calcium (**Supplementary Table 2**). Following the procedure outlined in **Fig. 1**, we obtained the activities for individual biosensors by averaging across cells with corresponding barcodes, identified by linear unmixing of spectral images. We stimulated cells with six pharmacological agents: 2-deoxyglucose (2DG), anisomycin, epidermal growth factor (EGF), ionomycin, phorbol-12,13-dibutyrate (PDBu), and UK14304, which were used to test these biosensors in previous studies (**Supplementary Table 2**). We analyzed the responses of all 24 biosensors to the agents and vehicle control (**Fig. 2** and **Supplementary Fig. 4-10**). As expected, the average biosensor activities responded to the known stimuli (**Fig. 2a, black boxes**). However, wide variations were noted between different cells expressing the same biosensor (**Fig. 2b**). Moreover, different biosensors that detect the same molecular activity may display distinct kinetics. For example, ERKKTR ^6^ and EKAR ^14^ both report ERK activation, but the kinetics of ERKKTR was delayed compared to that of EKAR, consistent with previous observations carried out in other cell types (**Fig. 2c**) ^15,16^.

**Figure 2.**
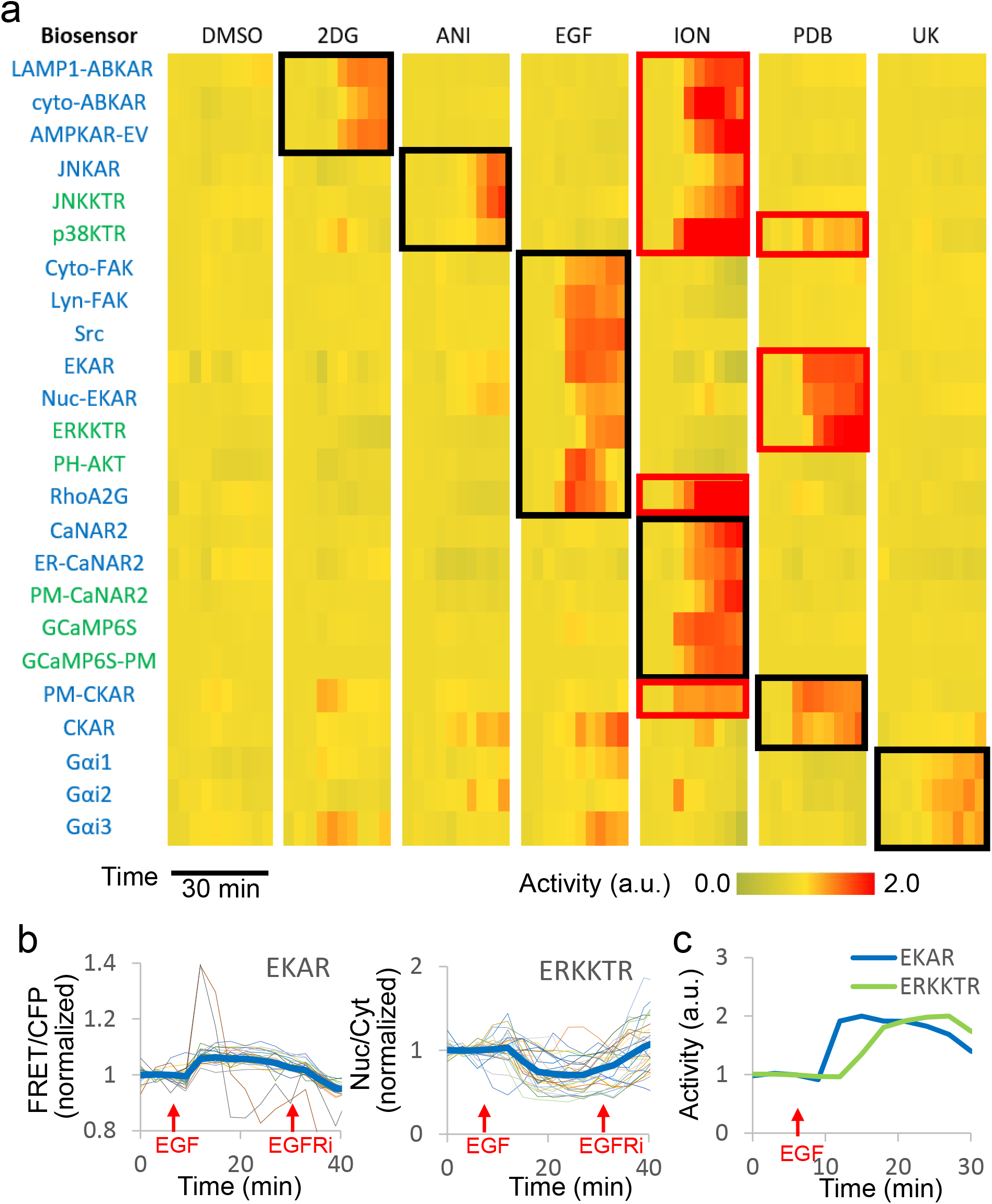
Multiplexed real-time monitoring of signaling activities by biosensor barcoding. Mixed population of barcoded HeLa cells expressing 24 different FRET (blue) and non-FRET (green) biosensors (see Supplementary Table 2) were stimulated with the indicated small molecule activators at 6 min. The activities normalized to pre-stimulus levels were averaged across cells of the corresponding barcode and adjusted for the dynamic range of each biosensor (see Methods). Black boxes indicate responses to known activators including: 2DG: 10 mM 2-deoxyglucose; ANI: 1 μg/ml anisomycin; EGF: 100 ng/ml EGF; ION: 1 μM ionomycin; PDB: 200 nM phorbol-12,13-dibutyrate; UK: 10 μM UK14304. Red boxes indicate additional responses (see text) (b) Traces of normalized FRET/CFP signal for EKAR (left) and nuclear-to-cytoplasm ratio of ERKKTR (right) for individual cells with the barcode corresponding to the two biosensors from a mixed population of cells. Thick blue lines represent the average responses. (c) Comparison of the kinetics of EKAR and ERKKTR in response to 100 ng/mL EGF obtained from mixed barcoded cells. Activities (normalized to the peak responses) represent average of n=46 (EKAR) and 22 (EKRKTR) cells.

To validate the results derived from mixed barcoded cells, we compared the responses to those from a homogeneous population of cells expressing single biosensors. The kinetics of the responses obtained from single cell populations were in general agreement with those from mixed cell populations (**Supplementary Fig. 11**). Additionally, the magnitude of the responses assessed at 9 and 15 min after stimulation showed no significant difference between the two groups for the majority (over 75%) of biosensors (**Supplementary Fig. 12**). Some of the differences noted between the two groups may be attributed to selection bias in analyzing single biosensor populations, in which only a small fraction of imaged cells—more likely the brighter ones—were selected, whereas in mixed biosensor populations all cells were included for analysis.

In addition to the expected activities, our analyses revealed responses to perturbations not included in the initial reports of the biosensors (**Fig. 2a, red boxes**), such as ionomycin-induced activation of AMPK, JNK, p38, RhoA, and PKC, as well PDBu-induced activation of ERK and p38. We validated these results using single population of HeLa cells expressing individual biosensors (**Supplementary Fig. 13**). Although several of these responses have been reported previously in different cell types using a variety of assays (**Supplementary Table 3**), our approach offers an efficient way to identify these events in the cells of interest and comprehensively characterize their kinetics. Note that the fluctuating activities of CKAR and Gαi1, 2, 3 in 2DG, anisomycin, and EGF experiments (**Fig. 2a**) were due to large cell-to-cell variation (**Supplementary Fig. 5-7**) relative to an overall small (~5%) dynamic range of these biosensors in our cells (**Supplementary Fig. 11**), rather than due to true responses to these perturbations. Taken together, these results demonstrate that: 1) the barcodes allow for correct identification of cells expressing different biosensors and are compatible with biosensor imaging without affecting biosensor responses; 2) the technique reveals variations between cells and kinetics of biosensors; and 3) multiplexed biosensors imaging can facilitate comprehensive identification and characterization of multiple biochemical responses to perturbations.

In conclusion, biosensor barcoding provides a simple method for simultaneously tracking large numbers of fluorescent biosensors. Given the ever-growing list of fluorescent biosensors, with a recent review listing over a thousand designs for monitoring ~170 cellular targets including proteins, voltage, ions, and metabolites^8^, we envision our method to find a wide range of applications, such as reconstructing the structure of complex molecular networks from timecourse perturbations, and the identification of cellular pathways targeted by pharmacological agents. In addition, biosensor barcoding can facilitate the development of new biosensors through side-by-side comparison of the sensitivity and dynamic range of multiple of biosensor designs. Newly developed biosensors will then further expand the biosensor barcoding technique. Although we only demonstrated the use of barcoding for CFP-YFP FRET and GFP based biosensors, our technique can be readily adapted to other types of fluorescent or bioluminescent biosensors with similar emission spectra.

## Methods

### Plasmids

#### Biosensors

Plasmids for biosensors were purchased from Addgene (see **Supplementary Table 2**). ERKKTR, p38KTR and JNKKTR genes were further cloned into pEGFP-N1 (CloneTech) via XhoI and BamHI restriction sites using the following primers: 5’-CAAgtcgacATGAAGGGCCGAAAGCCTC-3’ and 5’-CAAggatccccGGATGGGAATTGAAAGCTGGACT-3’ for ERKKTR, 5’-CAActcgagATGCGTAAGCCAGATCTCCG-3’ and 5’-CAAggatccccGCTGGACTGGAGGGTCAG-3’ for p38KTR; 5’-CAActcgagATGAGTAACCCTAAGATCCTAAAACAGAG-3’ and 5’-CAAggatccccGCTGGACTGGAGGGTCAG-3’ for JNKKTR.

#### Barcoding proteins

Barcoding proteins used in this study were derived from four fluorescent proteins (designated as A: TagRFP, B: mCardinal, C: iRFP702, and D: BFP) linked to targeting sequences of four subcellular locations (designated as 1: nucleus, 2: plasma membrane, 3: nuclear membrane, and 4: cytoplasm). Thus, a total of 16 barcoding proteins were generated, each represented by a letter-number combination (e.g. A1 indicates TagRFP targeted to the nucleus; see **Supplementary Fig. 2a**). To construct barcoding proteins, the following fragments were joined together by overlapping PCR: 1) fluorescent protein sequence; 2) spacer (TCTGGCAGCGGAGGCTCTGGAGGC); and 3) targeting sequence. The following plasmids were used as templates for the fluorescent proteins: H2B-TagRFP (Addgene #99271, a gift from Philipp Keller), mCardinal-H2B-C-10 (Addgene #56162)^11^, piRFP702-N1 (Addgene #45456)^12^, and EBFP2-Nucleus-7 (Addgene #55249, a gift from Michael Davidson). The targeting sequences for the nucleus, plasma membrane, nuclear membrane, and cytoplasm were derived from NLS of SV40, CAAX of K-Ras, lamin B1, and the NES of MAPKK, respectively. The complete sequences of the 16 barcoding proteins can be found in **Supplementary Note 1-16**.

We generated dual expression barcoding vectors by inserting pairs of barcoding protein sequences into the pVITRO1-hygro-mcs plasmid (InvivoGen), which contains two multiple cloning sites MCS1 and MCS2 (**Supplementary Fig. 2b**). The first barcoding protein sequence was inserted into MCS1 using BspEI and BamHI restriction sites. Subsequently, the second barcoding protein sequence was inserted into MCS2 using AgeI and BglII sites that are compatible with BspEI and BamHI, respectively (**Supplementary Fig. 2b**). As mentioned in the main text, the two barcoding proteins are of different colors and targeted to different subcellular locations, leading to a total of 72 possible combinations (**Supplementary Fig. 2c**).

### Chemical reagents

Stocks of 200 μM phorbol-12,13-dibutyrate (PDBu, EMD MilliporeSigma 524390), 1 mg/mL anisomycin (MilliporeSigma A9789), 10 mM UK14304 (MilliporeSigma U104), 10 mM yohimbine (MilliporeSigma Y3125), 1 mM gefitinib (Cayman 13166), and 1 mM ionomycin (Peprotech 5608212) were prepared by dissolving the chemicals in DMSO. Stocks were diluted to the indicated final concentrations in the culture medium. The EGF stock solution was prepared by dissolving EGF (MilliporeSigma E9644) in 10 mM acetic acid to a final concentration of 1 mg/ml. All drug stocks were stored at −20°C. 2-Deoxyglucose (2-DG, MilliporeSigma D8375) was dissolved in culture medium to 100 mM and used immediately.

### Cell lines

HeLa cells, purchased from ATCC, were grown at 37°C, 5% CO_2_ in DMEM high glucose medium (Gibco, #11965092) supplemented with 10% FBS (Corning Cellgro, 35-010-CV), 1 mM sodium pyruvate (Gibco, #11360070), and 1X non-essential amino acids (Gibco, #11140076). Transient transfections were performed using GenJet In Vitro DNA Transfection Reagent ver. II (SignaGen, #SL100499) following manufacturer’s instruction. Cells were transferred to 35 mm glass-bottom dishes (Mattek, Tissue Culture Dish P35GC-0-14-C) and allowed to attach overnight prior to imaging. For imaging of mixed population of barcoded cells expressing different biosensors, cells were mixed at equal ratios, seeded at 4×10^5^ per dish, and incubated at 37°C, 5% CO_2_ overnight before imaging experiments.

### Microscopy

Imaging experiments were carried out on a Zeiss LSM 780 or 880 single-point laser-scanning microscope (Zeiss AxioObserver with 780 or 880-Quasar confocal module; 34-channel spectral, high-sensitivity gallium arsenide phosphide (GaAsP) detectors) with a motorized stage for capturing multiple viewfields controlled by the Zen software as previously described ^17^. Live-cell imaging was carried out in a temperature/humidity/CO_2_-regulated chamber. To image barcodes, spectral images for red-far red barcoding proteins were acquired between 560 to 695 nm at 8.9 nm windows using Lambda Mode under 561 nm and 633 nm illumination. Reference spectra for TagRFP, mCardinal, and iRFP702 were acquired by imaging HeLa cells expressing H2B-TagRFP under 561 nm excitation, H2B-mCardinal under 561 and 633 nm excitation, and H2B-iRFP702 under 633 nm excitation. Since the BFP emission spectrum is well separated from those of the red-far red fluorophores (**Fig. 1a**), no unmixing is required for BFP images, which were therefore acquired in the Channel Mode. To avoid bleedthrough from CFP and YFP used in FRET-based biosensors, we collect BFP emission in the 370-430 nm range under 405 nm excitation (**Fig. 1a**). To image biosensors, CFP and YFP emissions under 457 nm illumination were obtained. This setting, while optimized for detecting CFP-YFP FRET biosensors, also captures GFP-based biosensors due to the overlapping spectra of GFP and YFP. Using a single imaging setting for both types of biosensors is convenient, and it reduces the cell exposure to illumination, therefore minimizing phototoxicity. Time-lapse images of the biosensors were taken at a rate of one frame every three minutes (unless specified), while the cells (in 2 ml DMEM, Gibco 21063029) were stimulated by adding signaling activators or inhibitors (volume: 200 μL for 2DG and 20 μL for all other reagents; concentration see above and **Fig. 2** legend) at the indicated time points.

### Image analysis

#### Analysis of barcodes by linear unmixing of spectral images

Using the Linear Unmixing function in the ZEN Software, spectra images of cells expressing pairs of barcoding proteins acquired under 561 nm illumination were unmixed using TagRFP and mCardinal reference spectra, whereas spectra images acquired under 633 um illumination were unmixed with mCardinal and iRFP702 spectra (**Supplementary Fig. 1**). The unmixed images for TagRFP, mCardinal, and iRFP702 as well as image of BFP were then combined in NIH ImageJ and Fiji ^18,19^ and visually inspected for the expression and localization of each fluorophore. In the majority of cases (>90%) the two expressed barcoding proteins can be unambiguously identified (see **Supplementary Fig. 3**). Cells with ambiguous barcodes were discarded for the analysis of biosensors. For simplicity, barcodes consist of four numbers that denote expression and location of TagRFP, mCardinal, iRFP702, and BFP, respectively (0: no expression; 1: nucleus; 2: plasma membrane, 3; nuclear membrane, and 4: cytoplasm; see **Fig. 1c**).

#### Analysis for biosensors in mixed populations of barcoded cells

Images of biosensors were processed and analyzed with NIH ImageJ and Fiji ^18,19^. The barcode of each cell, determined as described above, allows for identification of the biosensor expressed by the cell. To measure the activities for FRET based biosensors, the mean intensity of YFP over the entire cell was divided by that of CFP for each frame. For ERKKTR, p38KTR and JNKKTR, the mean intensity of YFP of the nucleus was divided by that of a cytoplasmic region. For PH-AKT, the mean intensity of YFP of an intracellular region was measured. For GCaMP6S and GCaMP6S-PM, the mean intensity of YFP over the entire cell was measured. The activities for every frame were then normalized to the average of those from the prestimulus frames. Normalized activities from cells with the same barcode were then pooled together to calculate the mean and standard deviation (S.D.) for the corresponding biosensor.

### Statistics

At least three independent experiments were carried out on different days for imaging experiments involving mixed barcoded cells (**Fig. 2, Supplementary Fig. 4-11**). Mean ± S.D. was reported as indicated in the figure legends. Statistics were derived by aggregating the number of samples noted in each figure legend across independent experiments. Statistical significance and p-values for comparing the responses of single and mixed cell populations (**Supplementary Fig. 12**) were determined using two-tailed unpaired Welch’s t-test for comparison between the two groups.

### Data availability

All data supporting the findings of the current study are available within the article and its Supplementary Information files or from the corresponding authors upon reasonable request.

## Acknowledgements

The authors would like to thank Hoku West-Foyle for helpful discussions. This work was supported by NIH grants K22CA212060 (to C.H.H.), R01GM136711 (to C.H.H.), Cervical Cancer SPORE P50CA098252 Career Development Award (to J.M.Y) and Pilot Project Award (to C.H.H.), and W. W. Smith Charitable Trust Cancer Research Grant #C1901 (to C.H.H.). The Zeiss LSM 780 and 880 confocal microscopes were purchased with NIH grants S10OD016374 and S10OD023548, respectively.

## Author contributions

J.M.Y. and C.H.H. conceived the project and designed the experiments; J.M.Y. and W.Y.C conducted the experiments; J.M.Y., W.Y.C.,J.L., and C.H.H. analyzed the data; J.M.Y. and C.H.H. wrote the manuscript with inputs from W.Y.C. and J.L; C.H.H. and J.M.Y. supervised the study.

