## Supplementary Information for "Highly multiplexed imaging of biosensors in live cells"

### Supplementary Figure 1

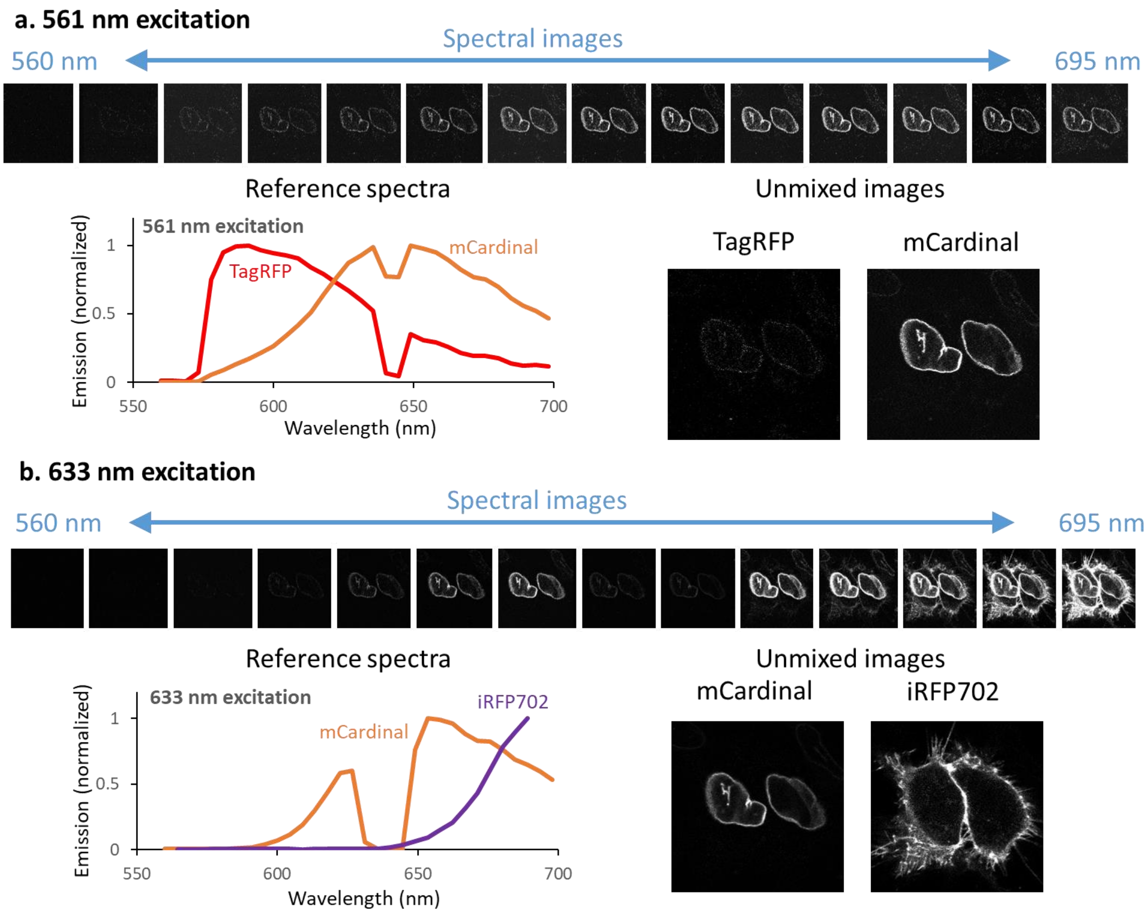


**Supplementary Figure 1. Unmixing the barcoding protein signals.** HeLa cells expressing TagRFP, mCardinal, or iRFP702 targeted to different subcellular locations were spectrally imaged on a Zeiss LSM 780 confocal microscope. The images were decomposed into the component colors by linear unmixing of the spectral images using the reference spectra obtained from cells expressing individual fluorophores under the same laser wavelength of excitation. Specifically, spectral images acquired under 561 nm excitation were unmixed into TagRFP and mCardinal (a), whereas those under 633 nm excitation were unmixed into mCardinal and iRFP702 (b). In the example shown, the cells expressed nuclear membrane-targeted mCardinal as well as plasma membrane targeted iRFP702. Note that the dips in the spectra were due to blocking of emission by the dichroic mirror used for reflecting the excitation beam.

### Supplementary Figure 2

a

| | A. RFP | B. mCar | C. iRFP | D. BFP |
|---|---|---|---|---|
| 1. Nuc | A1 | B1 | C1 | D1 |
| 2. PM | A2 | B2 | C2 | D2 |
| 3. NM | A3 | B3 | C3 | D3 |
| 4. Cyt | A4 | B4 | C4 | D4 |

b

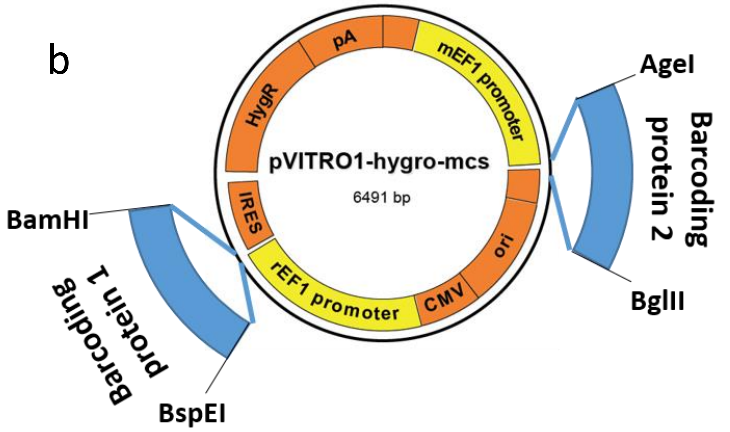


c

| Barcode | MCS1 | MCS2 |
|---|---|---|
| 1200 | A1 | B2 |
| 1300 | A1 | B3 |
| 1400 | A1 | B4 |
| 1020 | A1 | C2 |
| 1030 | A1 | C3 |
| 1040 | A1 | C4 |
| 1002 | A1 | D2 |
| 1003 | A1 | D3 |
| 1004 | A1 | D4 |
| 2100 | A2 | B1 |
| 2300 | A2 | B3 |
| 2400 | A2 | B4 |
| 2010 | A2 | C1 |
| 2030 | A2 | C3 |
| 2040 | A2 | C4 |
| 2001 | A2 | D1 |
| 2003 | A2 | D3 |
| 2004 | A2 | D4 |
| 3100 | A3 | B1 |
| 3200 | A3 | B2 |
| 3400 | A3 | B4 |
| 3010 | A3 | C1 |
| 3020 | A3 | C2 |
| 3040 | A3 | C4 |

| Barcode | MCS1 | MCS2 |
|---|---|---|
| 3001 | A3 | D1 |
| 3002 | A3 | D2 |
| 3004 | A3 | D4 |
| 4100 | A4 | B1 |
| 4200 | A4 | B2 |
| 4300 | A4 | B3 |
| 4010 | A4 | C1 |
| 4020 | A4 | C2 |
| 4030 | A4 | C3 |
| 4001 | A4 | D1 |
| 4002 | A4 | D2 |
| 4003 | A4 | D3 |
| 0120 | B1 | C2 |
| 0130 | B1 | C3 |
| 0140 | B1 | C4 |
| 0102 | B1 | D2 |
| 0103 | B1 | D3 |
| 0104 | B1 | D4 |
| 0210 | B2 | C1 |
| 0230 | B2 | C3 |
| 0240 | B2 | C4 |
| 0201 | B2 | D1 |
| 0203 | B2 | D3 |
| 0204 | B2 | D4 |

| Barcode | MCS1 | MCS2 |
|---|---|---|
| 0310 | B3 | C1 |
| 0320 | B3 | C2 |
| 0340 | B3 | C4 |
| 0301 | B3 | D1 |
| 0302 | B3 | D2 |
| 0304 | B3 | D4 |
| 0410 | B4 | C1 |
| 0420 | B4 | C2 |
| 0430 | B4 | C3 |
| 0401 | B4 | D1 |
| 0402 | B4 | D2 |
| 0403 | B4 | D3 |
| 0012 | C1 | D2 |
| 0013 | C1 | D3 |
| 0014 | C1 | D4 |
| 0021 | C2 | D1 |
| 0023 | C2 | D3 |
| 0024 | C2 | D4 |
| 0031 | C3 | D1 |
| 0032 | C3 | D2 |
| 0034 | C3 | D4 |
| 0041 | C4 | D1 |
| 0042 | C4 | D2 |
| 0043 | C4 | D3 |

**Supplementary Figure 2. Dual expression barcoding vectors.** (a) Barcoding proteins generated from combinations of fluorescent proteins and targeting sequences. (b) Map of pVITRO-hygro-mcs vector used for generating dual expressing barcoding vectors. Coding sequences of a pair of barcoding proteins were cloned into the two MCS sites of the pVITRO1 vector. (c) 72 dual expression barcoding vectors generated from pairs of barcoding proteins inserted into the two MCS sites of pVITRO-hygro-mcs (see Fig. 1c for barcode nomenclature).

#### Supplementary Figure 3

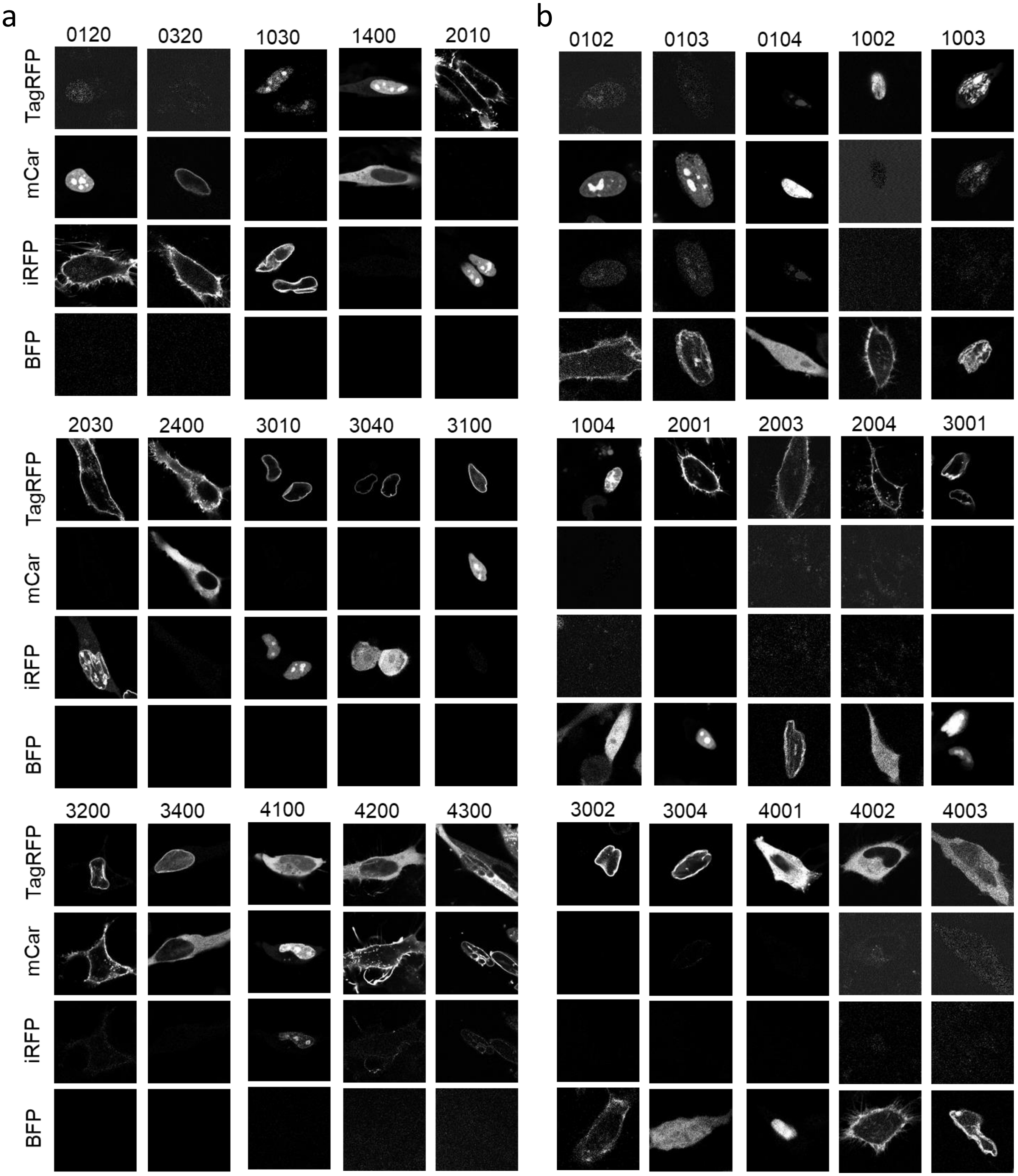


**Supplementary Figure 3. Barcodes generated from four FPs.** 30 out of 72 possible barcodes are shown. (a) Examples of HeLa cells expressing pairs of red/far-red barcoding proteins. (b) Examples of HeLa cells expressing BFP and one of the three red/far-red FPs. Spectral images of cells expressing pairs of barcoding proteins targeted to different subcellular locations were unmixed to obtain three images corresponding to TagRFP, mCardinal, and iRFP702, respectively. BFP is imaged separately. The numbers on top of every set of images denote barcodes as defined in Fig. 1c legend.

### Supplementary Figure 4 (DMSO)

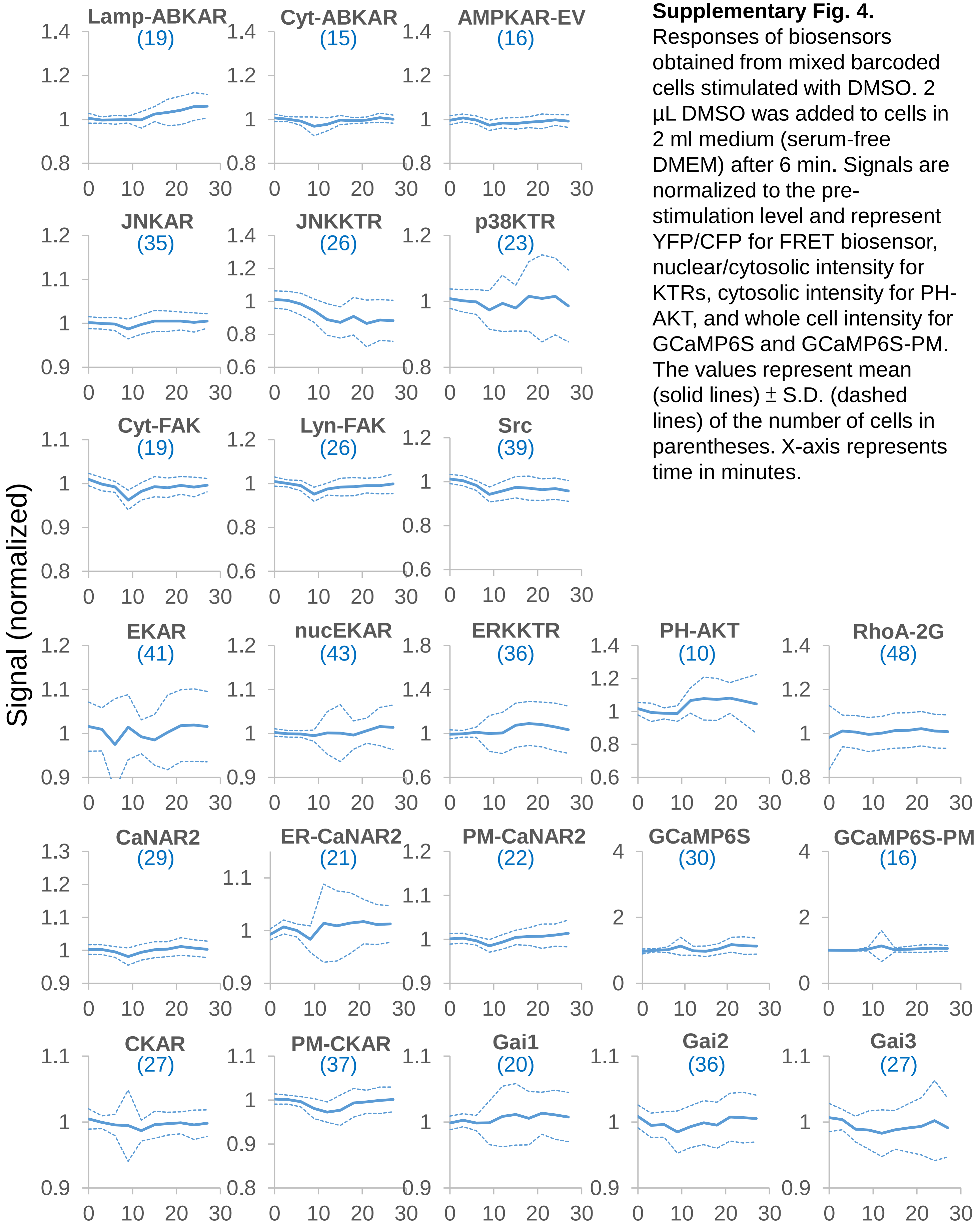


**Supplementary Fig. 4.** Responses of biosensors obtained from mixed barcoded cells stimulated with DMSO. 2 µL DMSO was added to cells in 2 ml medium (serum-free DMEM) after 6 min. Signals are normalized to the pre-stimulation level and represent YFP/CFP for FRET biosensor, nuclear/cytosolic intensity for KTRs, cytosolic intensity for PH-AKT, and whole cell intensity for GCaMP6S and GCaMP6S-PM. The values represent mean (solid lines) ± S.D. (dashed lines) of the number of cells in parentheses. X-axis represents time in minutes.

### Supplementary Figure 5 (2DG)

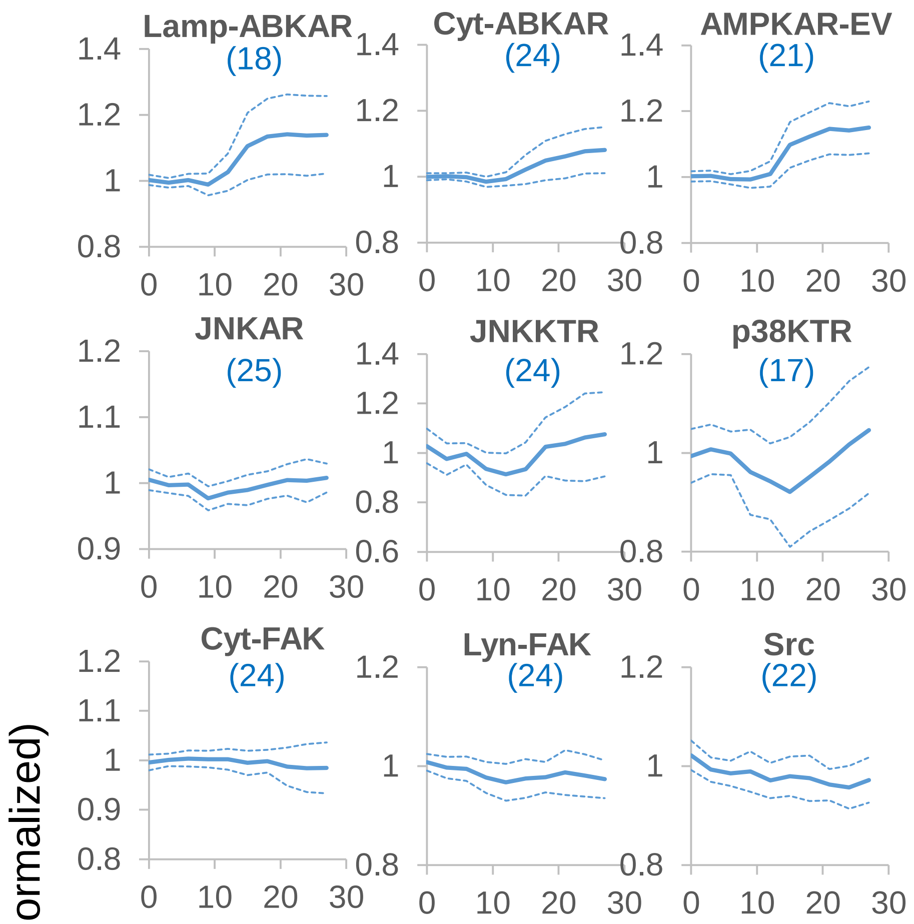


**Supplementary Fig. 5.** Responses of biosensors obtained from mixed barcoded cells stimulated with 2DG. 10 mM 2-deoxyglucose was given after 6 min. The signals are normalized to the pre-stimulation level and represent YFP/CFP for FRET biosensor, nuclear/cytosolic intensity for KTRs, cytosolic intensity for PH-AKT, and whole cell intensity for GCaMP6S and GCaMP6S-PM. The values represent mean (solid lines) ± S.D. (dashed lines) of the number of cells in parentheses. X-axis represents time in minutes.

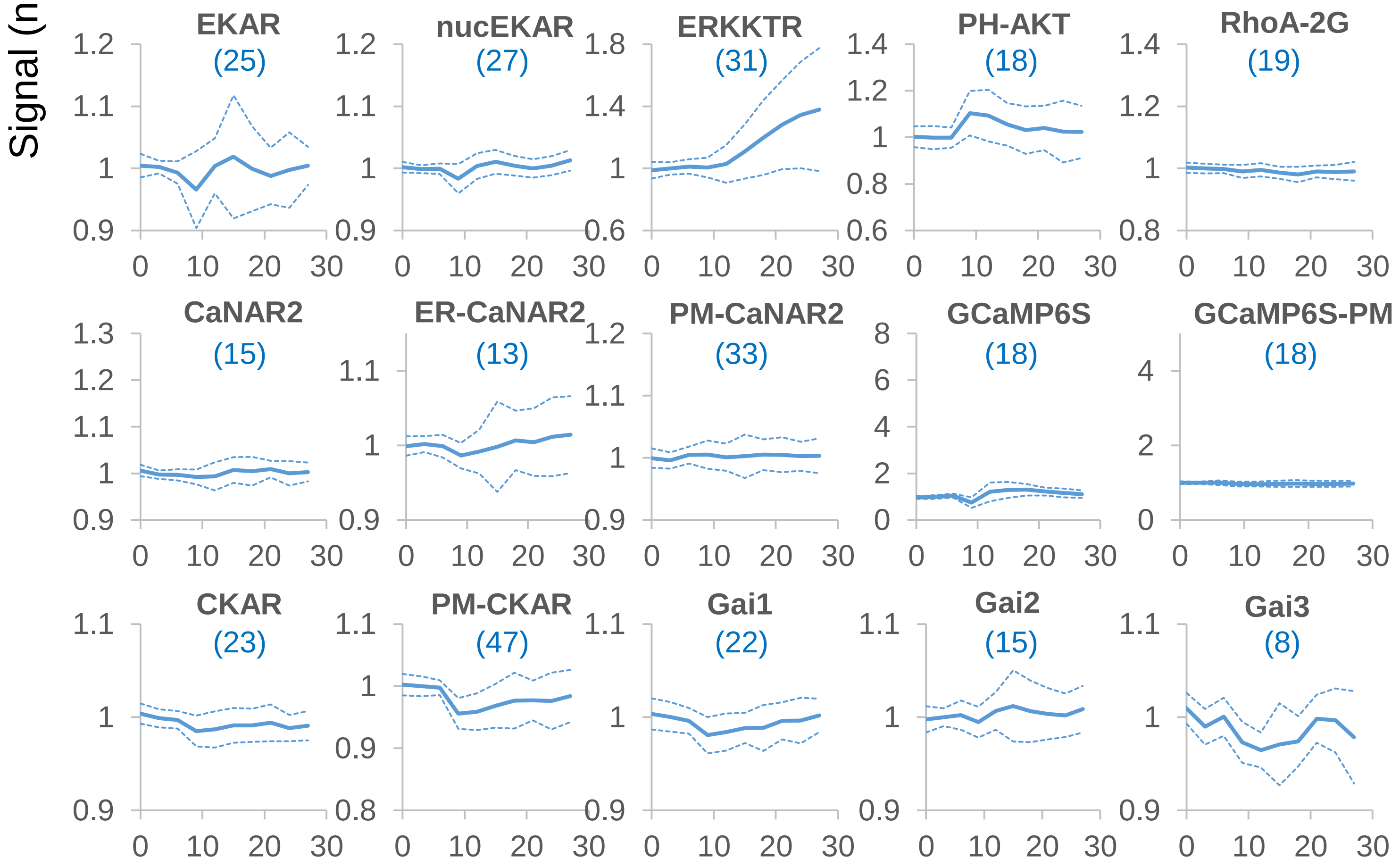

### Supplementary Figure 6 (Anisomycin)

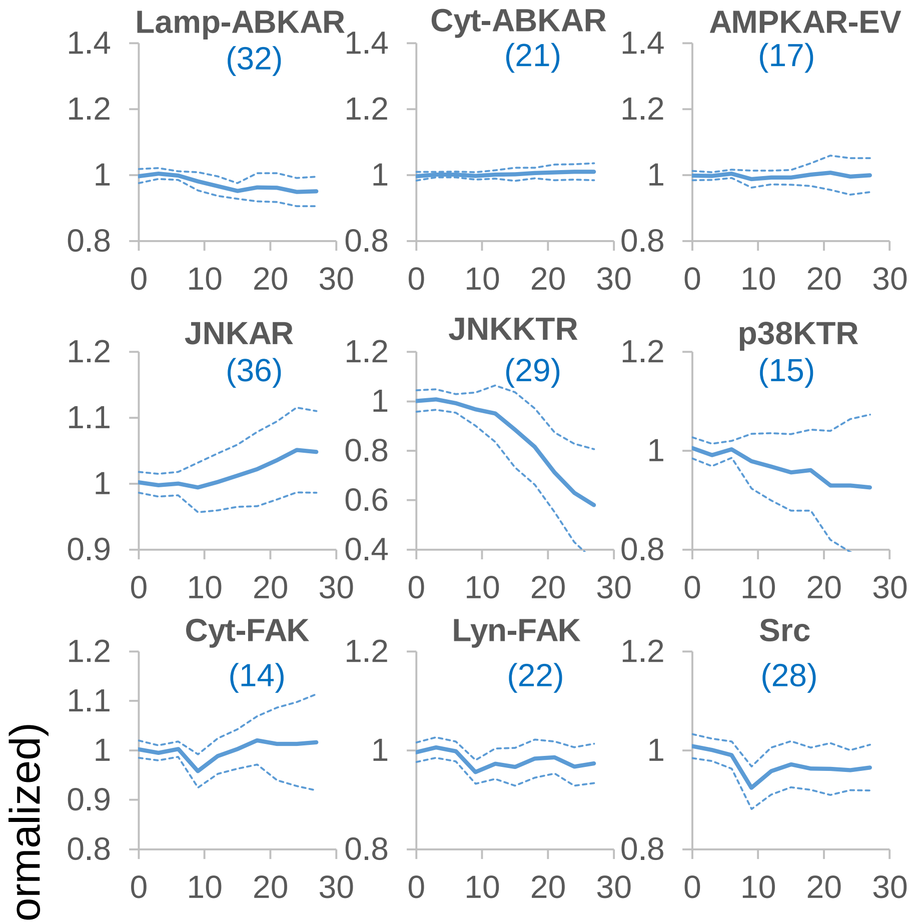


**Supplementary Fig. 6.** Responses of biosensors obtained from mixed barcoded cells stimulated with anisomycin. 1 µg/ml anisomycin was given after 6 min. The signals are normalized to the pre-stimulation level and represent YFP/CFP for FRET biosensor, nuclear/cytosolic intensity for KTRs, cytosolic intensity for PH-AKT, and whole cell intensity for GCaMP6S and GCaMP6S-PM. The values represent mean (solid lines) ± S.D. (dashed lines) of the number of cells in parentheses. X-axis represents time in minutes.

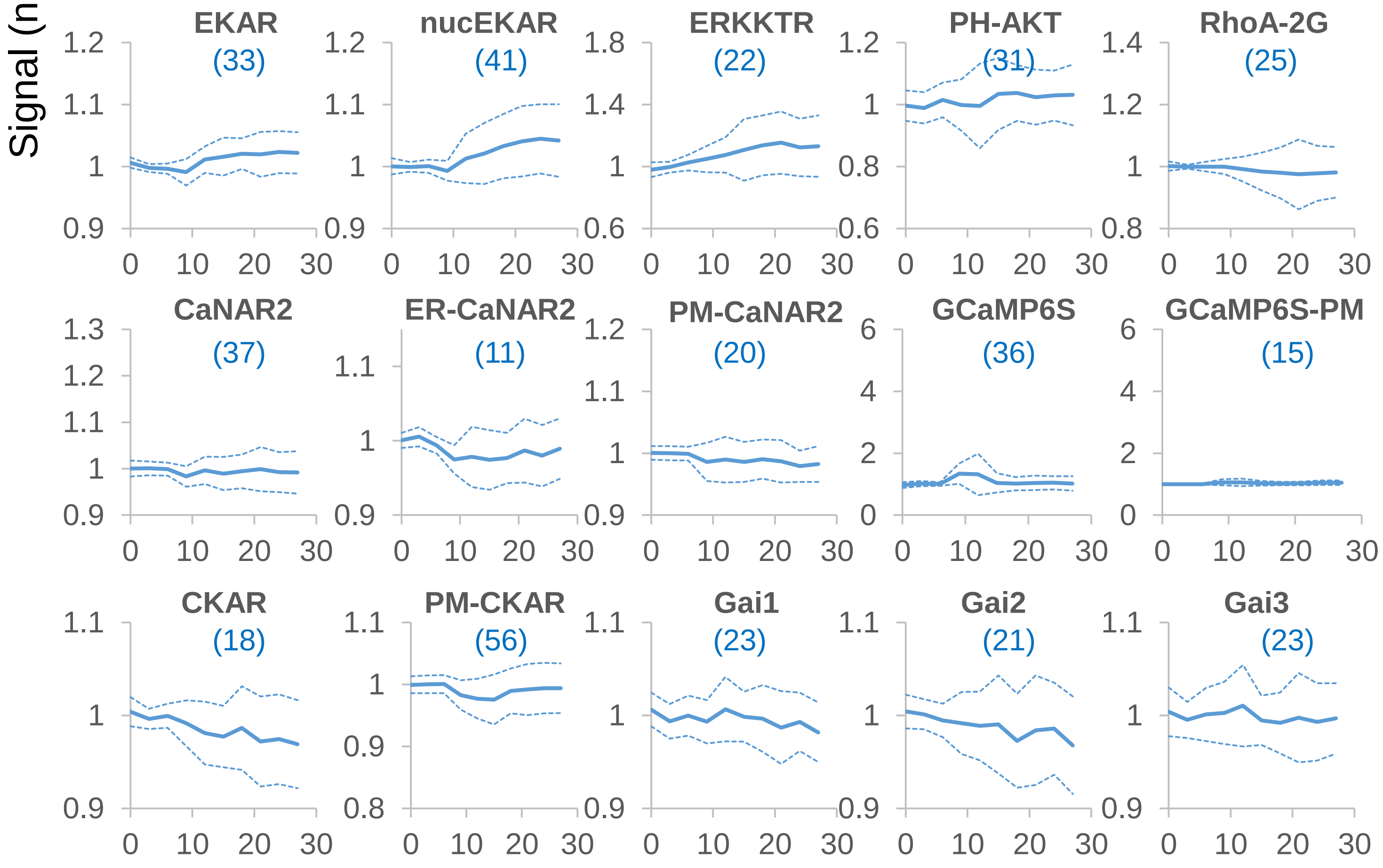

### Supplementary Figure 7 (EGF)

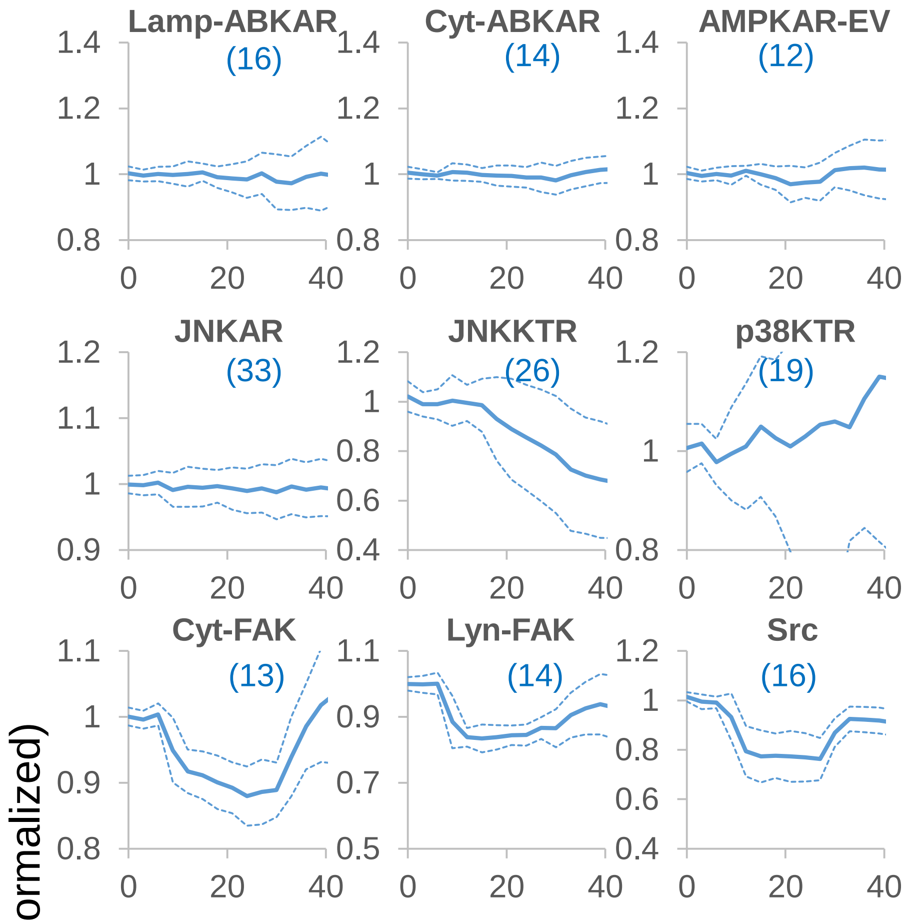


**Supplementary Fig. 7.** Responses of biosensors obtained from mixed barcoded cells stimulated with EGF and EGFRi. 100 ng/ml EGF was given after 6 min; 1 µM gefinitib was given at 27 min. The signals are normalized to the pre-stimulation level and represent YFP/CFP for FRET biosensor, nuclear/cytosolic intensity for KTRs, cytosolic intensity for PH-AKT, and whole cell intensity for GCaMP6S and GCaMP6S-PM. The values represent mean (solid lines) ± S.D. (dashed lines) of the number of cells in parentheses. X-axis represents time in minutes.

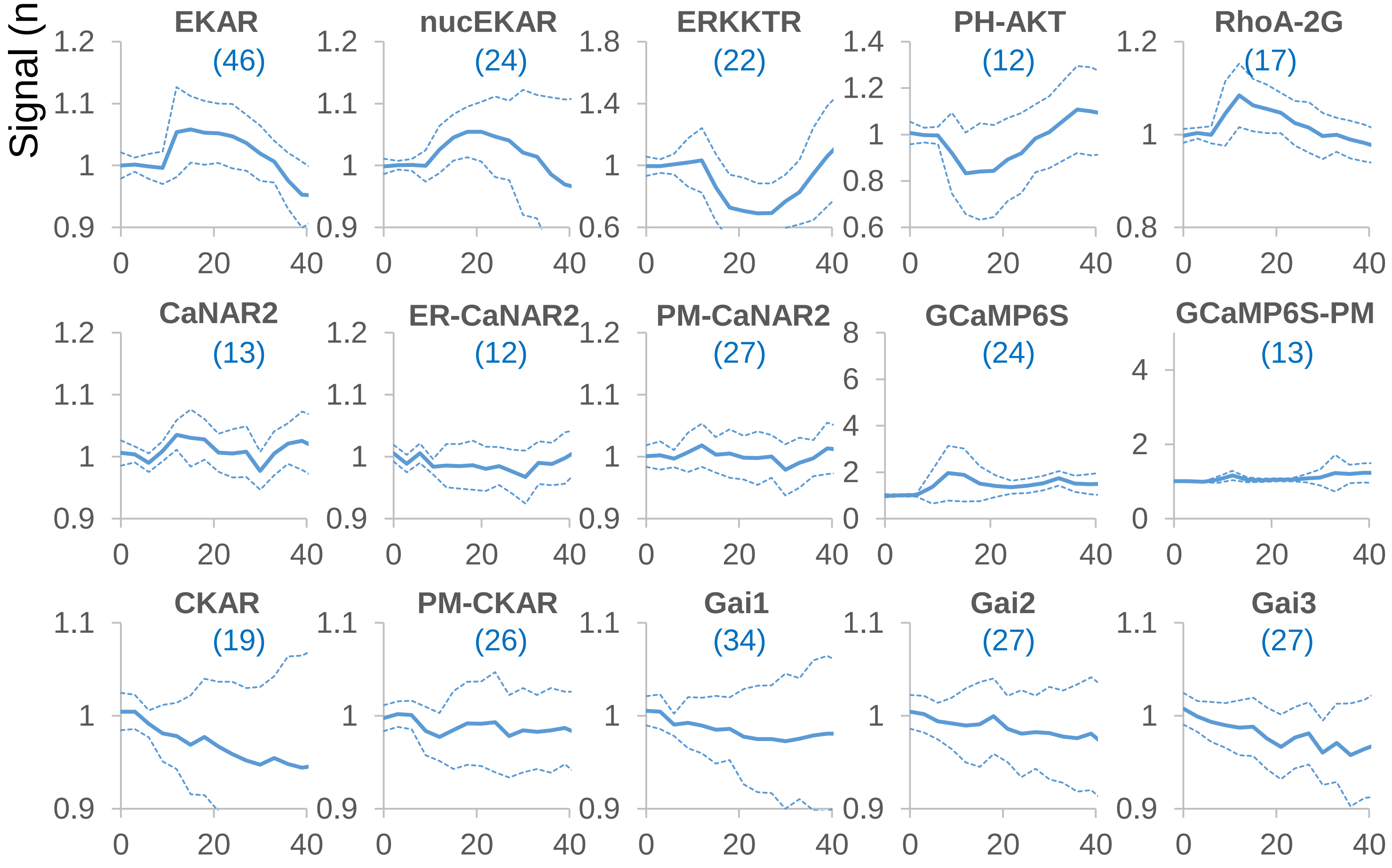

### Supplementary Figure 8 (Ionomycin)

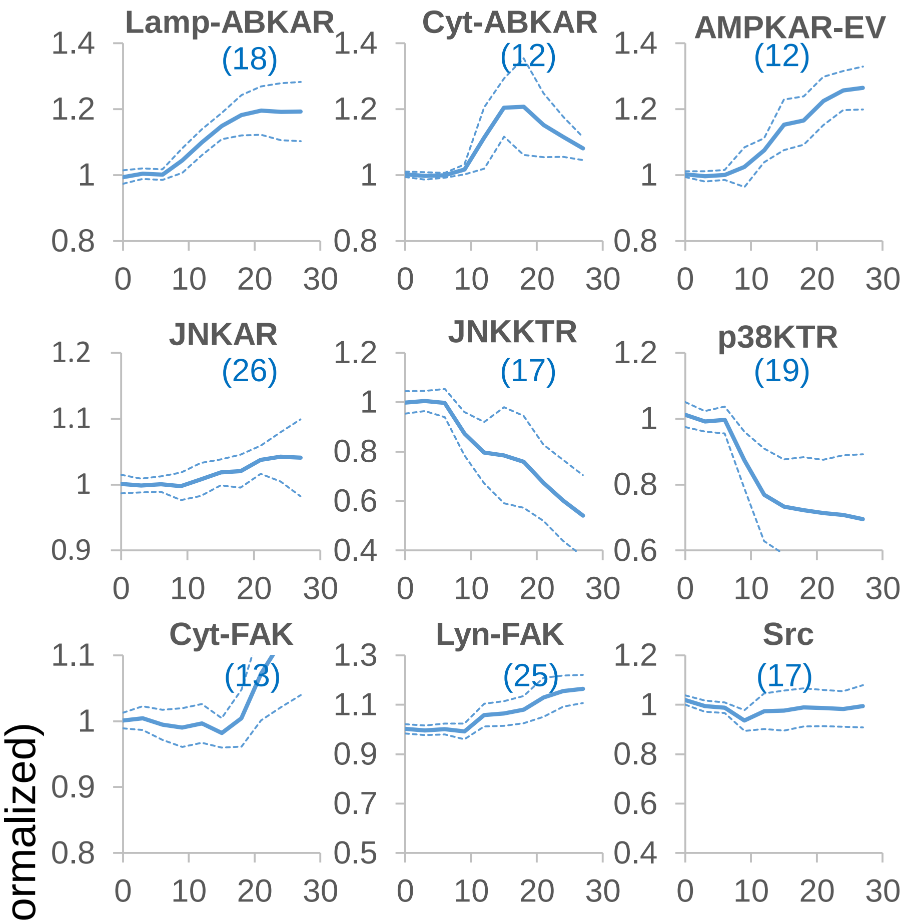


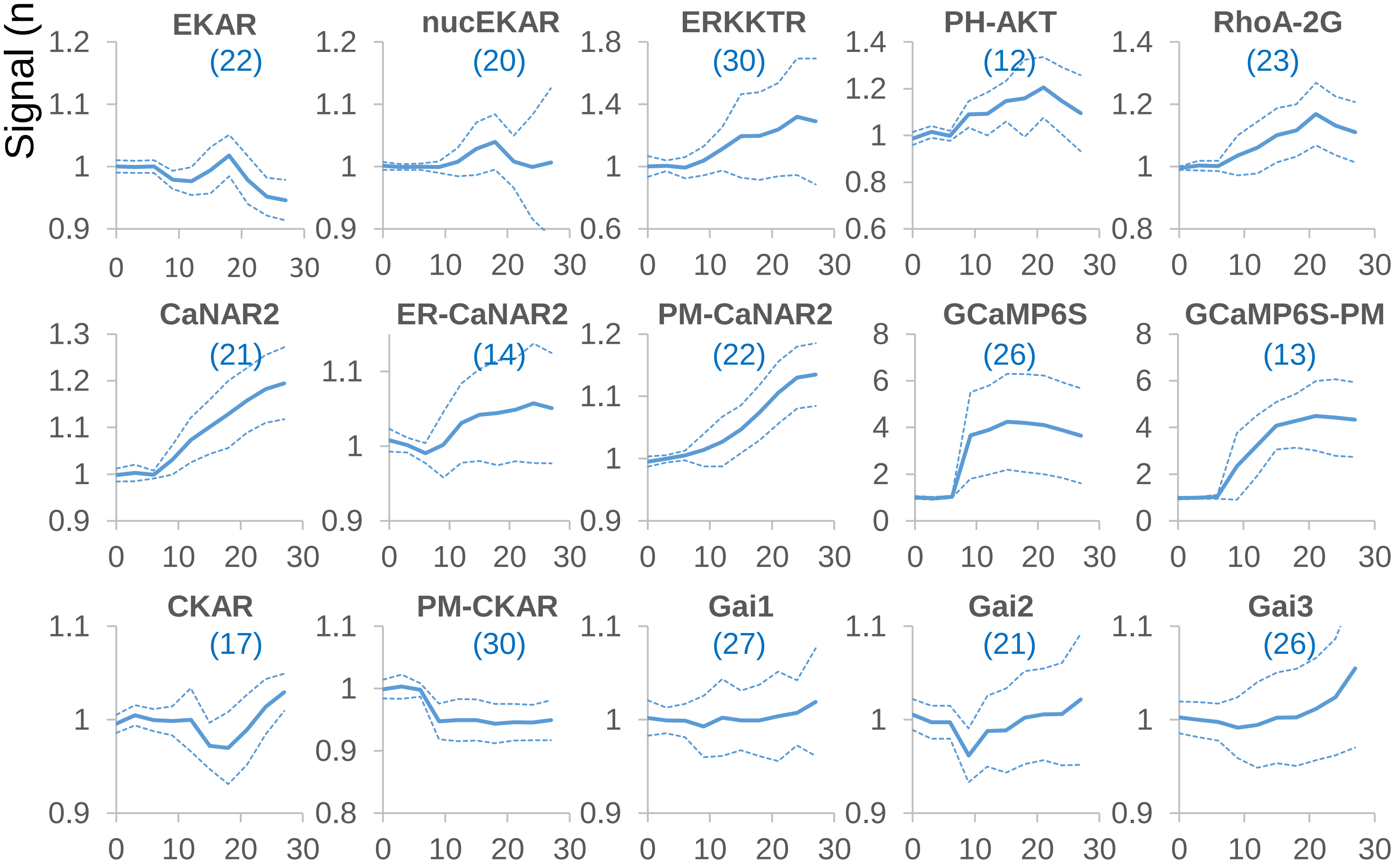


**Supplementary Fig. 8.** Responses of biosensors obtained from mixed barcoded cells stimulated with ionomycin. 1 µM ionomycin was given after 6 min. The signals are normalized to the pre-stimulation level and represent YFP/CFP for FRET biosensor, nuclear/cytosolic intensity for KTRs, cytosolic intensity for PH-AKT, and whole cell intensity for GCaMP6S and GCaMP6S-PM. The values represent mean (solid lines) ± S.D. (dashed lines) of the number of cells in parentheses. X-axis represents time in minutes.

### Supplementary Figure 9 (PDBu)

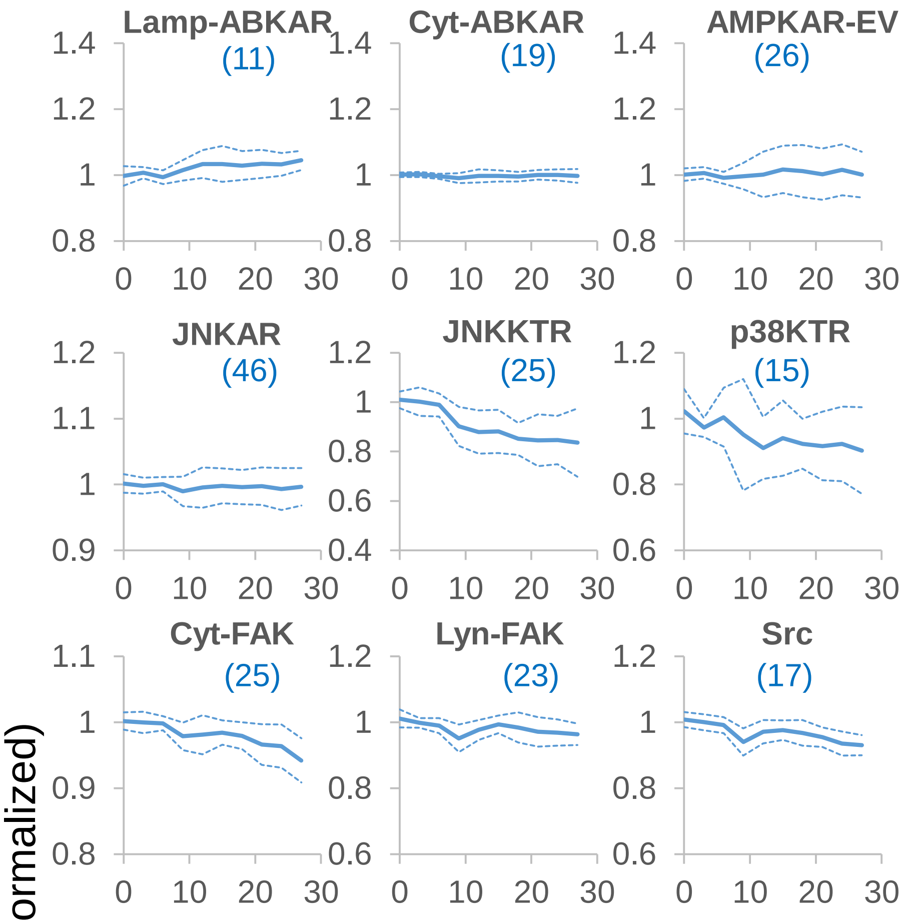


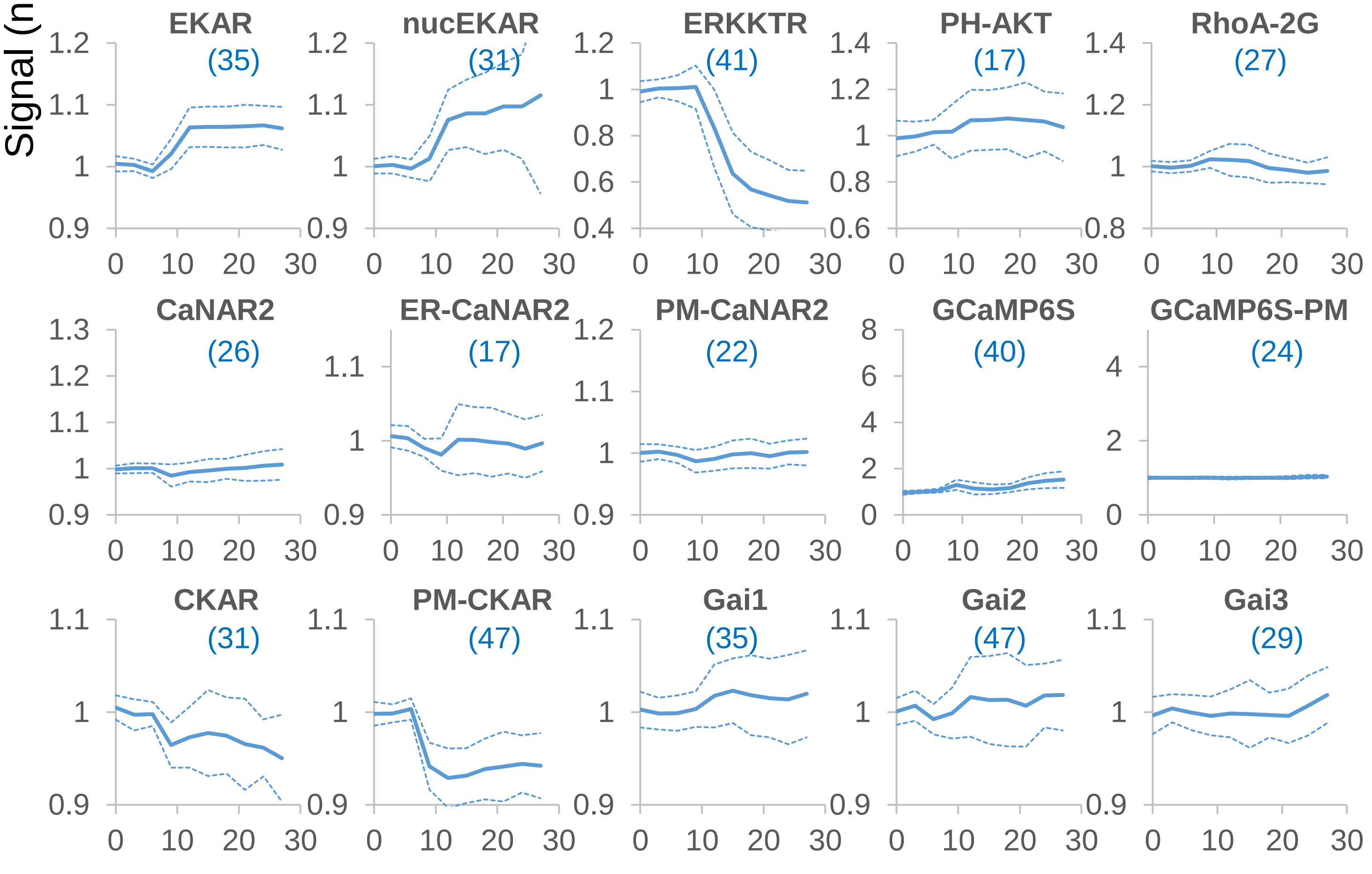


**Supplementary Fig. 9.** Responses of biosensors obtained from mixed barcoded cells stimulated with PDBu (phorbol-12,13-dibutyrate). 200 nM PDBu was given after 6 min. The signals are normalized to the pre-stimulation level and represent YFP/CFP for FRET biosensor, nuclear/cytosolic intensity for KTRs, cytosolic intensity for PH-AKT, and whole cell intensity for GCaMP6S and GCaMP6S-PM. The values represent mean (solid lines) ± S.D. (dashed lines) of the number of cells in parentheses. . X-axis represents time in minutes.

### Supplementary Figure 10 (UK14304)

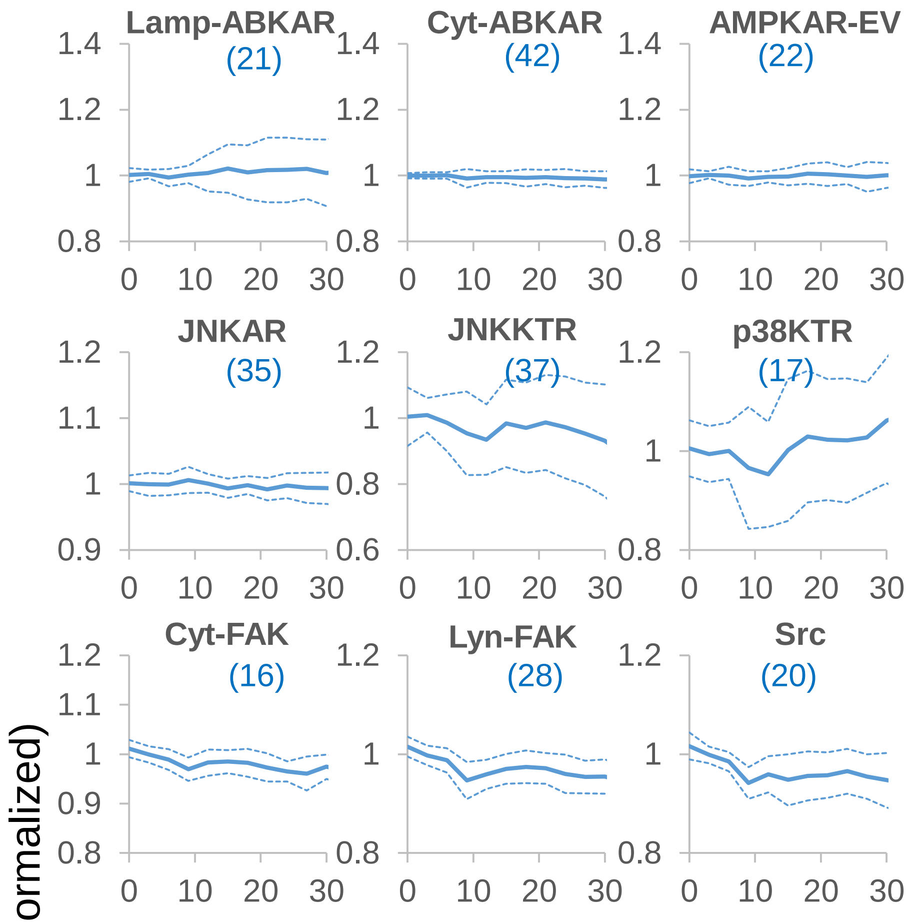


**Supplementary Fig. 10.** Responses of biosensors obtained from mixed barcoded cells stimulated with UK14304. 10 µM UK14304 was given after 6 min. The signals are normalized to the pre-stimulation level and represent YFP/CFP for FRET biosensor, nuclear/cytosolic intensity for KTRs, cytosolic intensity for PH-AKT, and whole cell intensity for GCaMP6S and GCaMP6S-PM. The values represent mean (solid lines) ± S.D. (dashed lines) of the number of cells in parentheses. X-axis represents time in minutes.

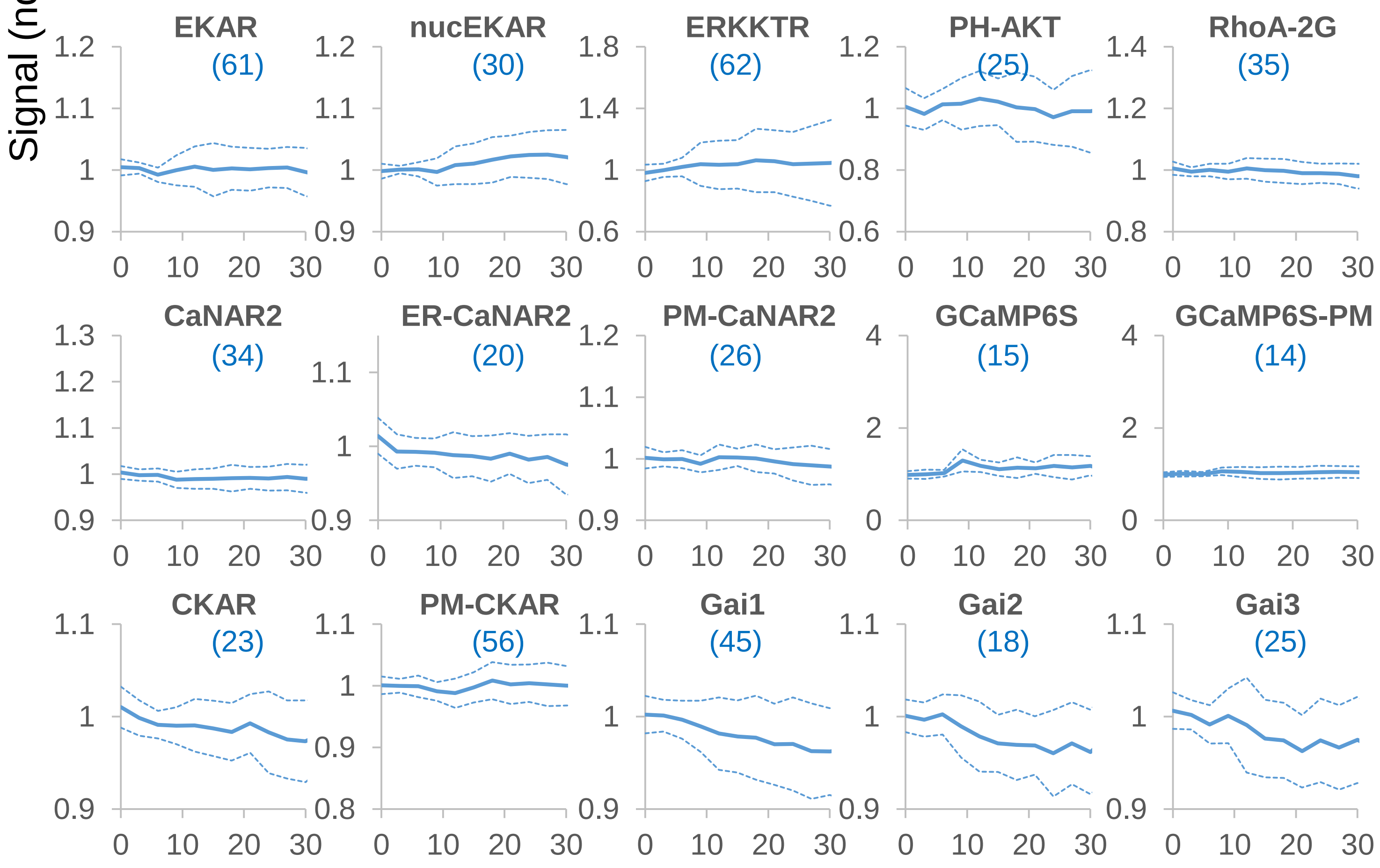

### Supplementary Figure 11

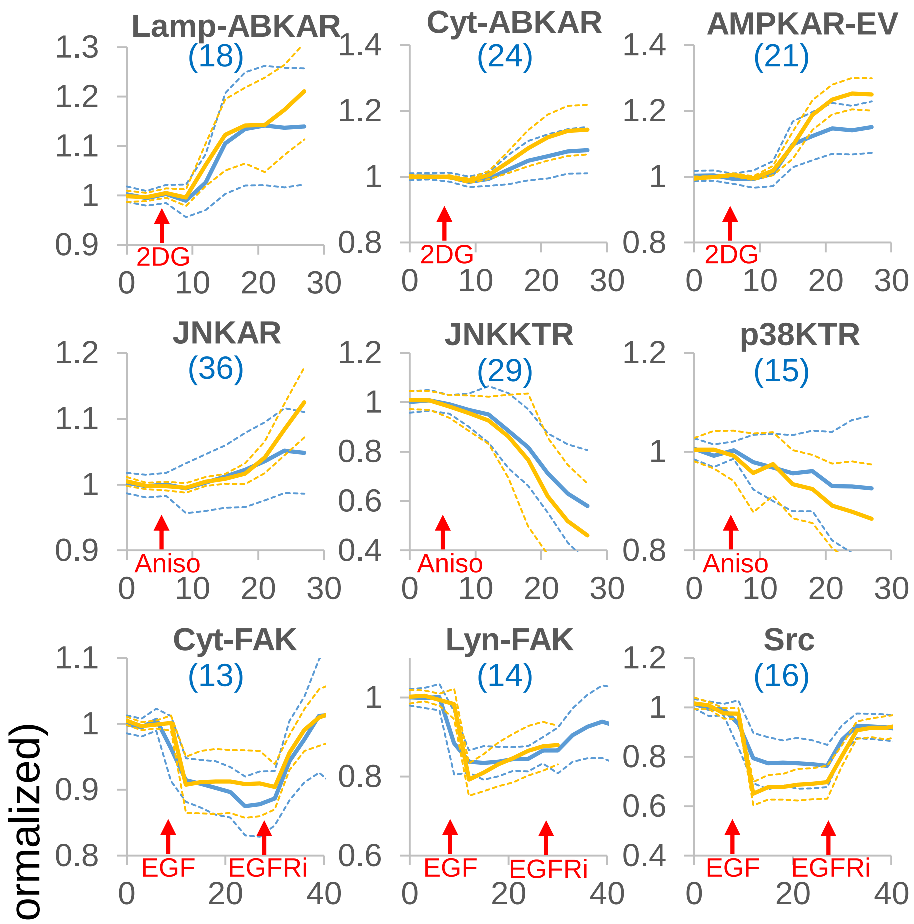


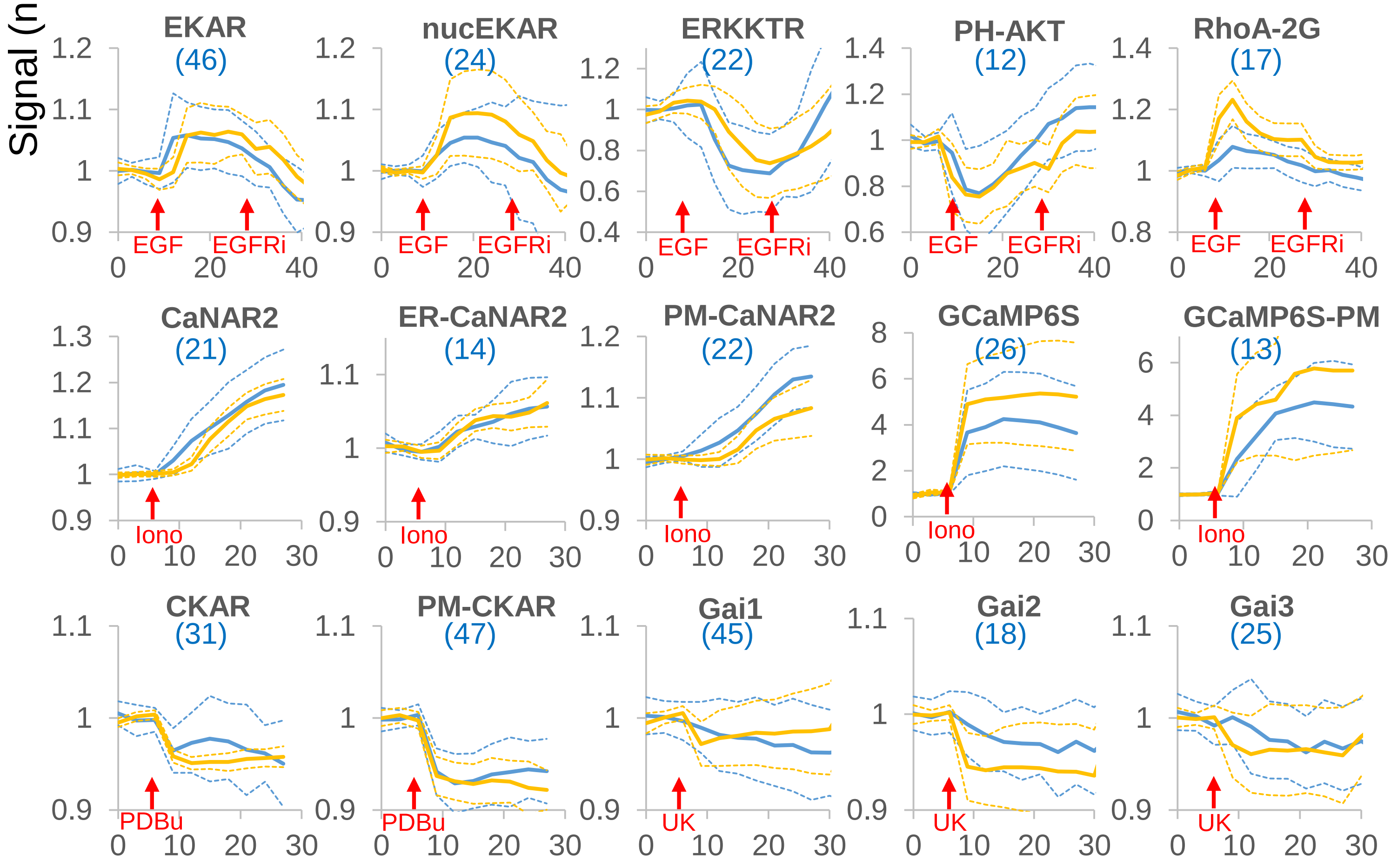


**Supplementary Fig. 11.** Comparison of the responses of biosensors obtained from mixed barcoded cells (blue) and single cell population (orange). Activators including 2DG (10 mM 2-deoxyglucose), Aniso (1 µg/ml anisomycin), EGF (100 ng/ml EGF); Iono (1 µM ionomycin), PDBu (200 nM phorbol-12,13-dibutyrate) and UK (10 µM UK14304) were given after 6 min. EGFRi (1 µM gefitinib) was given after 27 min. The signals are normalized to the pre-stimulation level and represent YFP/CFP for FRET biosensor, nuclear/cytosolic intensity for KTRs, cytosolic intensity for PH-AKT, and whole cell intensity for GCaMP6S and GCaMP6S-PM. The values represent mean (solid lines) ± S.D. (dashed lines). n=20 cells for single cell population and shown in parenthesis for mixed barcoded cells. X-axis represents time in minutes.

### Supplementary Figure 12

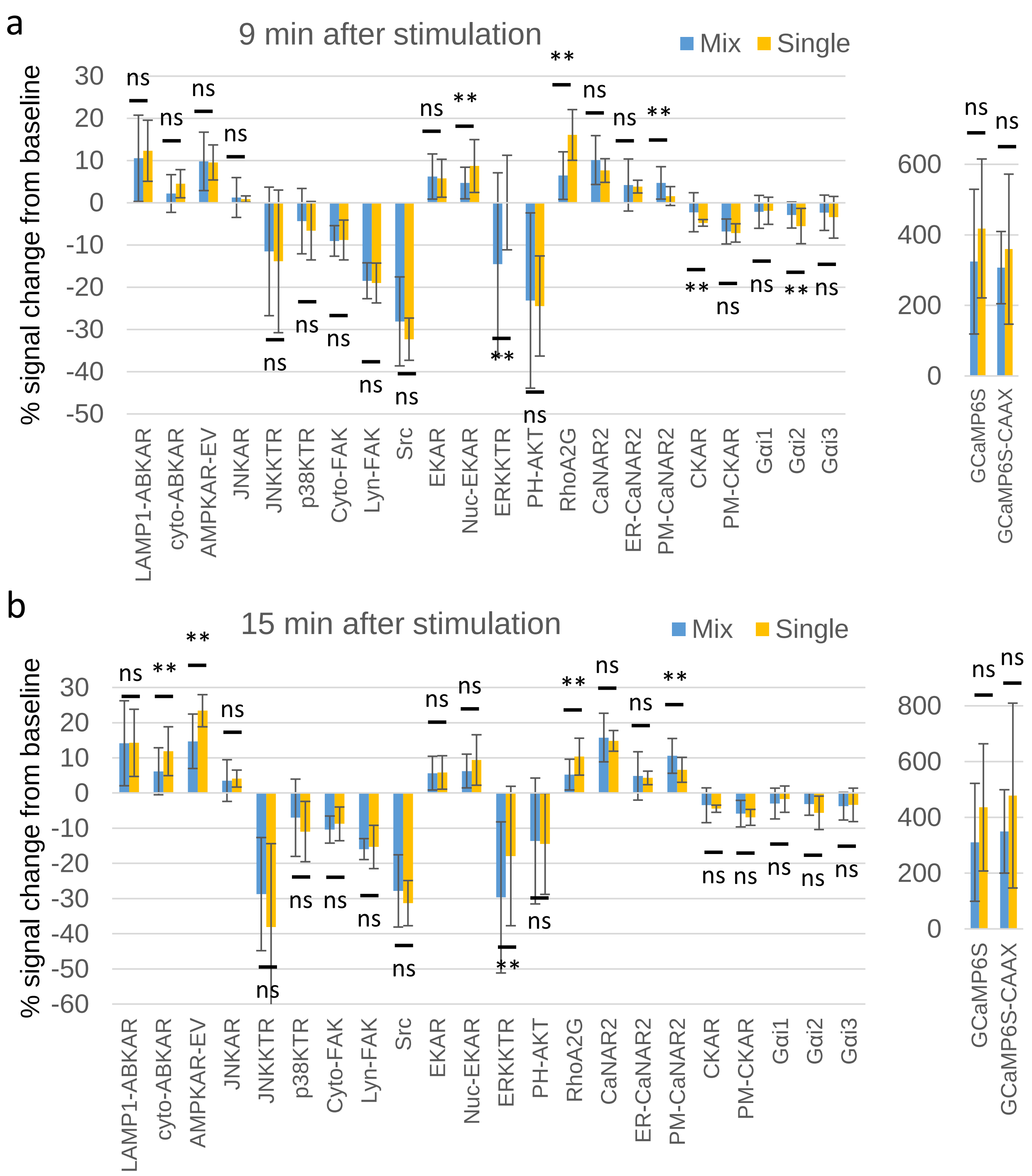


**Supplementary Fig. 12.** Comparison of the magnitude of responses at 9 min (a) and 15 min (b) after stimulation for biosensors obtained from mixed barcoded cells (blue) and single cell population (orange). See Supplementary Fig. 11 for the activators used as well as the definition of signal for each biosensor. Statistical significance and p-values for the two groups were determined using two-tailed unpaired Welch's t-test. Error bars: S.D. ns: no significant difference; **: p<0.05.

### Supplementary Figure 13

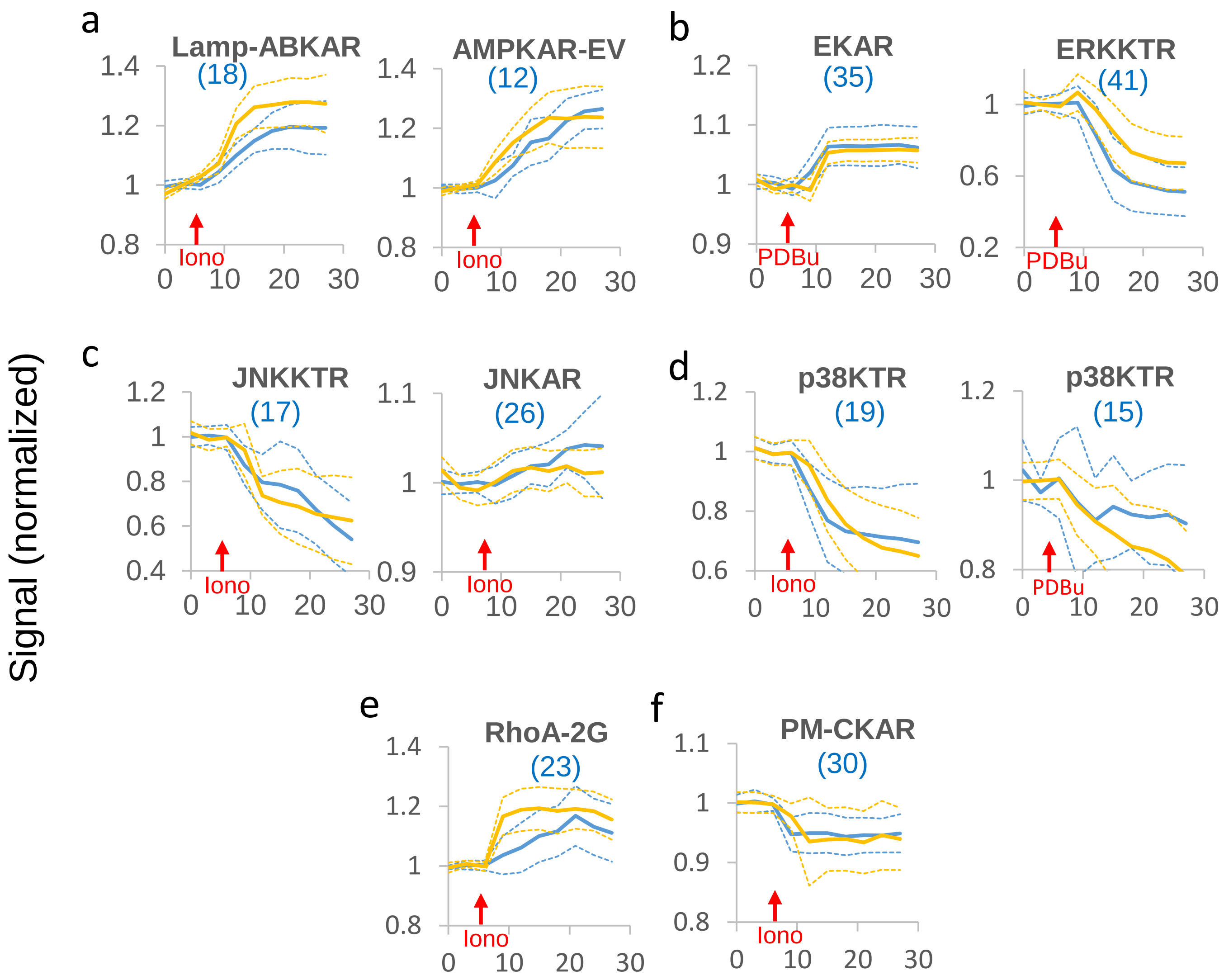


**Supplementary Fig. 13. Validation of additional biosensor responses identified in mixed populations of barcoded cells.** Responses of biosensors to 1 µM ionomycin (iono) or 200 nM PDBu obtained from mixed barcoded cells (blue) are plotted with those from single cell populations (orange) expressing biosensors for (a) AMPK; (b) ERK; (c) JNK; (d) p38; (e) RhoA; and (f) PKC. The signals are normalized to the pre-stimulation level and represent YFP/CFP for FRET biosensor and nuclear/cytosolic intensity for KTRs. The values represent mean (solid lines) ± S.D. (dashed lines). n=20 cells for single cell population and shown in parenthesis for mixed barcoded cells. X-axis represents time in minutes.

**Supplementary Table 1.** Theoretical number of barcodes generated from pairs of barcoding proteins of different fluorophores and targeted to different subcellular locations. Red numbers correspond to the current study.

| # targeting sites \ # fluorophores | 3 | 4 | 5 | 6 |
|---|---|---|---|---|
| 3 | 18 | 36 | 60 | 90 |
| 4 | 36 | 72 | 120 | 180 |
| 5 | 60 | 120 | 200 | 300 |
| 6 | 90 | 180 | 300 | 450 |
| 7 | 126 | 252 | 420 | 630 |

**Supplementary Table 2.** List of targets and biosensors used in barcoded cells

| Target | Activator | Biosensor | Site* | Barcode | Source | Ref |
|---|---|---|---|---|---|---|
| AMPK | 2DG | AMPKAR-EV | | 1040 | Addgene #105241 | 21 |
| | | cyto-ABKAR | C | 1002 | Addgene #61510 | 22 |
| | | LAMP1-ABKAR | L | 0320 | Addgene #65068 | 22 |
| Calcineurin | Ionomycin | CaNAR2 | | 4020 | Addgene #64728 | 23 |
| | | ER-CaNAR2 | ER | 0120 | Addgene #64732 | 23 |
| | | PM-CaNAR2 | PM | 3040 | Addgene #64730 | 23 |
| Calcium | Ionomycin | GCaMP6S | | 4100 | Addgene #40753 | 24 |
| | | GCaMP6S-PM | PM | 4200 | Addgene #52228 | 25 |
| ERK | EGF | EKAR | C | 2100 | Addgene #18679 | 14 |
| | | Nuc-EKAR | N | 1300 | Addgene #18681 | 14 |
| | | ERKKTR | | 4001 | Addgene #59150 | 6 |
| FAK | EGF | Cyto-FAK | | 3002 | Addgene #78300 | 26 |
| | | Lyn-FAK | PM | 1020 | Addgene #78299 | 26 |
| Galphai1 | UK14034 | Gαi1 | | 2004 | Addgene #69623 | 27 |
| Galphai2 | UK14034 | Gαi2 | | 2040 | Addgene #69624 | 27 |
| Galphai3 | UK14034 | Gαi3 | | 3100 | Addgene #69625 | 27 |
| JNK | Anisomycin | JNKAR | | 2300 | Addgene #61625 | 28 |
| | | JNKKTR | | 2001 | Addgene #59151 | 6 |
| p38 | Anisomycin | p38KTR | | 3001 | Addgene #59152 | 6 |
| PI3K | EGF | PH-AKT | | 1004 | | 29 |
| PKC | PDBu | CKAR | | 3004 | Addgene #14860 | 30 |
| | | PM-CKAR | PM | 0102 | Addgene #14862 | 30 |
| RhoA | EGF | RhoA2G | | 4300 | Addgene #40176 | 31 |
| Src | EGF | Src | PM | 1200 | Addgene #78302 | 32 |

*C: cytosol; L: lysosome; ER: endoplasmic reticulum; PM: plasma membrane; N: nucleus

**Supplementary Table 3.** Reported responses of biosensor targets

| Target | Biosensor/Assay | Stimulation | Cells | Ref |
|---|---|---|---|---|
| AMPK | ABKAR | Ionomycin | hippocampal neuron | 33 |
| ERK | EKAR, nucEKAR | PDBu | HEK293 | 34 |
| p38 | Phospho-p38 immunoblot | Ionomycin | Neutrophil | 35 |
| JNK | Phospho-JNK immunoblot | Ionomycin | Jurkat | 36 |
| RhoA | 35S-Met labeling, Immunoblot | Ionomycin | intestinal epithelial cells | 37 |
| PKC | 32P labeling | Ionomycin | T cell | 38 |
| p38 | Phospho-p38 immunoblot | PDBu | cardiomyocytes, endometrial cells | 39, 40 |

#### Supplementary Note 1. Sequence of barcoding protein A1 (TagRFP-NLS)

| | |
|---|---|
| **ATGGTGTCA…** | **TagRFP** |
| **TCTGGCAGC…** | **Spacer** |
| **GATCCAAAA…** | **SV40 NLS 3x repeat** |
| **TGA** | **Stop** |

**ATGGTGTCAAAAGGTGAAGAACTGATTAAGGAAAATATGCACATGAAGTTATACATGGAAGGCACGGTC**
**AATAATCACCACTTTAAATGTACATCTGAAGGAGAGGGAAAACCGTACGAAGGAACTCAGACTATGCGA**
**ATCAAAGTCGTGGAGGGTGGCCCGCTGCCTTTCGCCTTCGATATTTTGGCTACTAGCTTCATGTACGGA**
**TCTCGCACCTTTATTAACCATACCCAGGGAATCCCAGATTTCTTTAAACAATCGTTCCCGGAAGGTTTC**
**ACGTGGGAAAGAGTGACGACATACGAAGATGGAGGAGTCCTCACCGCTACCCAGGACACGAGTTTGCAA**
**GATGGATGCTTGATCTATGATGTAAAGATTCGAGGCGTAAATTTCCCTTCGAACGGCCCAGTCATGCAA**
**AAGAAGACTTTGGGATGGGAAGCAAATACTGAAATGCTTTATCCTGCTGATGGAGGTCTTGAAGGACGT**
**ACGGATATGGCACTGAAACTGGTGGGCGGAGGACACTTGATATGTAATTTCAAAACTACGTATCGAAGT**
**AAAAAGCCGGCCAAGAATCTGAAAATGCCAGGAGTTTATTACGTTGACCACAGACTTGAAAGAATTAAG**
**GAGGCAGATAAAGAAACGTACGTGGAACAACACGAAGTAGCCGTTGCTCGCTATTGTGACTTGCCATCG**
**AAACTTGGACATAAACTTAATGGAATGGATGAACTTTATAAATCTGGCAGCGGAGGCTCTGGAGGCgga**
**agctccGATCCAAAAAAGAAGAGAAAGGTAGACCCCAAAAAGAAAAGGAAGGTGGACCCTAAGAAAAAG**
**CGCAAAGTGTGA**

#### Supplementary Note 2. Sequence of barcoding protein B1 (mCardinal-NLS)

| | |
|---|---|
| ATGGTGAGC… | mCardinal |
| TCTGGCAGC… | Spacer |
| GATCCAAAA… | SV40 NLS 3x repeat |
| TGA | Stop |

ATGGTGAGCAAGGGCGAGGAGCTGATCAAGGAGAACATGCACATGAAGCTGTACATGGAAGGCACCGTG
AACAACCACCACTTCAAGTGCACCACCGAAGGGGAGGGCAAGCCCTACGAGGGCACCCAGACCCAGAGG
ATTAAGGTGGTGGAGGGAGGCCCCCTGCCGTTCGCATTCGACATCCTGGCCACCTGCTTTATGTACGGG
AGCAAGACCTTCATCAACCACACCCAGGGCATCCCCGATTTCTTTAAGCAGTCCTTCCCTGAGGGCTTC
ACATGGGAGAGAGTCACCACATACGAAGACGGGGGCGTGCTTACCGTTACCCAGGACACCAGCCTCCAG
GACGGCTGCTTGATCTACAACGTCAAGCTCAGAGGGGTGAACTTCCCATCCAACGGCCCTGTGATGCAG
AAGAAAACACTCGGCTGGGAGGCCACCACCGAGACCCTGTACCCCGCTGACGGCGGCCTGGAAGGCAGA
TGCGACATGGCCCTGAAGCTCGTGGGCGGGGGCCACCTGCACTGCAACCTGAAGACCACATACAGATCC
AAGAAACCCGCTAAGAACCTCAAGATGCCCGGCGTCTACTTTGTGGACCGCAGACTGGAAAGAATCAAG
GAGGCCGACAATGAGACCTACGTCGAGCAGCACGAGGTGGCTGTGGCCAGATACTGCGACCTCCCTAGC
AAACTGGGGCACAAACTTAATGGCATGGACGAGCTGTACAAGTCTGGCAGCGGAGGCTCTGGAGGCgga
agctccGATCCAAAAAAGAAGAGAAAGGTAGACCCCAAAAAGAAAAGGAAGGTGGACCCTAAGAAAAAG
CGCAAAGTGTGA

#### Supplementary Note 3. Sequence of barcoding protein C1 (iRFP702-NLS)

| | |
|---|---|
| ATGGCGCGT… | iRFP702 |
| TCTGGCAGC… | Spacer |
| GATCCAAAA… | SV40 NLS 3x repeat |
| TGA | Stop |

ATGGCGCGTAAGGTCGATCTCACCTCCTGCGATCGCGAGCCGATCCACATCCCCGGCAGCATTCAGCCG
TGCGGCTGCCTGCTAGCCTGCGACGCGCAGGCGGTGCGGATCACGCGCATTACGGAAAATGCCGGCGCG
TTCTTTGGACGCGAAACTCCGCGGGTCGGTGAGCTACTCGCCGATTACTTCGGCGAGACCGAAGCCCAT
GCGCTGCGCAACGCACTGGCGCAGTCCTCCGATCCAAAGCGACCGGCGCTGATCTTCGGTTGGCGCGAC
GGCCTGACCGGCCGCACCTTCGACATCTCGCTGCATCGCCATGACGGTACATCGATCATCGAGTTCGAG
CCTGCGGCGGCCGAACAGGCCGACAATCCGCTGCGGCTGACGCGGCAGATCATCGCGCGCACCAAAGAA
CTGAAGTCGCTCGAAGAGATGGCCGCACGGGTGCCGCGCTATCTGCAGGCGATGCTCGGCTATCACCGC
GTGATGTTGTACCGCTTCGCGGACGACGGCTCCGGCAAAGTGATCGGCGAGGCGAAGCGCAGCGACCTC
GAGAGCTTTCTCGGTCAGCACTTTCCGGCGTCGCTGGTCCCGCAGCAGGCGCGGCTACTGTACTTGAAG
AACGCGATCCGCGTGGTCTCGGATTCGCGCGGCATCAGCAGCCGGATCGTGCCCGAGCACGACGCCTCC
GGCGCCGCGCTTGATCTGTCGTTCGCGCACCTGCGCAGCATCTCGCCTATCCATCTCGAATTTCTGCGG
AACATGGGCGTCAGCGCCTCGATGTCGCTGTCGATCATCATTGACGGCACGCTATGGGGATTGATCATC
TGTCATCATTACGAGCCGCGTGCCGTGCCGATGGCGCAGCGCGTCGCGGCCGAAATGTTCGCCGACTTC
TTATCGCTGCACTTCACCGCCGCCCACCACCAACGCTCTGGCAGCGGAGGCTCTGGAGGCggaagctcc
GATCCAAAAAAGAAGAGAAAGGTAGACCCCAAAAAGAAAAGGAAGGTGGACCCTAAGAAAAAGCGCAAA
GTGTGA

#### Supplementary Note 4. Sequence of barcoding protein D1 (BFP-NLS)

| | |
|---|---|
| ATGGTGAGC… | BFP |
| TCTGGCAGC… | Spacer |
| GATCCAAAA… | SV40 NLS 3x repeat |
| TGA | Stop |

ATGGTGAGCAAGGGCGAGGAGCTGTTCACCGGGGTGGTGCCCATCCTGGTCGAGCTGGACGGCGACGTA
AACGGCCACAAGTTCAGCGTGAGGGGCGAGGGCGAGGGCGATGCCACCAACGGCAAGCTGACCCTGAAG
TTCATCTGCACCACCGGCAAGCTGCCCGTGCCCTGGCCCACCCTCGTGACCACCCTGAGCCACGGCGTG
CAGTGCTTCGCCCGCTACCCCGACCACATGAAGCAGCACGACTTCTTCAAGTCCGCCATGCCCGAAGGC
TACGTCCAGGAGCGCACCATCTTCTTCAAGGACGACGGCACCTACAAGACCCGCGCCGAGGTGAAGTTC
GAGGGCGACACCCTGGTGAACCGCATCGAGCTGAAGGGCGTCGACTTCAAGGAGGACGGCAACATCCTG
GGGCACAAGCTGGAGTACAACTTCAACAGCCACAACATCTATATCATGGCCGTCAAGCAGAAGAACGGC
ATCAAGGTGAACTTCAAGATCCGCCACAACGTGGAGGACGGCAGCGTGCAGCTCGCCGACCACTACCAG
CAGAACACCCCCATCGGCGACGGCCCCGTGCTGCTGCCCGACAGCCACTACCTGAGCACCCAGTCCGTG
CTGAGCAAAGACCCCAACGAGAAGCGCGATCACATGGTCCTGCTGGAGTTCCGCACCGCCGCCGGGATC
ACTCTCGGCATGGACGAGCTGTACAAGTCTGGCAGCGGAGGCTCTGGAGGCggaagctccGATCCAAAA
AAGAAGAGAAAGGTAGACCCCAAAAAGAAAAGGAAGGTGGACCCTAAGAAAAAGCGCAAAGTGTGA

#### Supplementary Note 5. Sequence of barcoding protein A2 (TagRFP-CAAX)

| | |
|---|---|
| ATGGTGTCA… | TagRFP |
| TCTGGCAGC… | Spacer |
| AAAGAAAAG… | CAAX |
| TGA | Stop |

ATGGTGTCAAAAGGTGAAGAACTGATTAAGGAAAATATGCACATGAAGTTATACATGGAAGGCACGGTC
AATAATCACCACTTTAAATGTACATCTGAAGGAGAGGGAAAACCGTACGAAGGAACTCAGACTATGCGA
ATCAAAGTCGTGGAGGGTGGCCCGCTGCCTTTCGCCTTCGATATTTTGGCTACTAGCTTCATGTACGGA
TCTCGCACCTTTATTAACCATACCCAGGGAATCCCAGATTTCTTTAAACAATCGTTCCCGGAAGGTTTC
ACGTGGGAAAGAGTGACGACATACGAAGATGGAGGAGTCCTCACCGCTACCCAGGACACGAGTTTGCAA
GATGGATGCTTGATCTATGATGTAAAGATTCGAGGCGTAAATTTCCCTTCGAACGGCCCAGTCATGCAA
AAGAAGACTTTGGGATGGGAAGCAAATACTGAAATGCTTTATCCTGCTGATGGAGGTCTTGAAGGACGT
ACGGATATGGCACTGAAACTGGTGGGCGGAGGACACTTGATATGTAATTTCAAAACTACGTATCGAAGT
AAAAAGCCGGCCAAGAATCTGAAAATGCCAGGAGTTTATTACGTTGACCACAGACTTGAAAGAATTAAG
GAGGCAGATAAAGAAACGTACGTGGAACAACACGAAGTAGCCGTTGCTCGCTATTGTGACTTGCCATCG
AAACTTGGACATAAACTTAATGGAATGGATGAACTTTATAAATCTGGCAGCGGAGGCTCTGGAGGCAAA
GAAAAGATGAGCAAAGATGGTAAAAAGAAGAAAAAGAAGTCAAAGACAAAGTGTGTAATTATGTGA

#### Supplementary Note 6. Sequence of barcoding protein B2 (mCardinal-CAAX)

**ATGGTGAGC…** **mCardinal**
**TCTGGCAGC…** **Spacer**
**AAAGAAAAG…** **CAAX**
**TGA** **Stop**

**ATGGTGAGCAAGGGCGAGGAGCTGATCAAGGAGAACATGCACATGAAGCTGTACATGGAAGGCACCGTG**
**AACAACCACCACTTCAAGTGCACCACCGAAGGGGAGGGCAAGCCCTACGAGGGCACCCAGACCCAGAGG**
**ATTAAGGTGGTGGAGGGAGGCCCCCTGCCGTTCGCATTCGACATCCTGGCCACCTGCTTTATGTACGGG**
**AGCAAGACCTTCATCAACCACACCCAGGGCATCCCCGATTTCTTTAAGCAGTCCTTCCCTGAGGGCTTC**
**ACATGGGAGAGAGTCACCACATACGAAGACGGGGGCGTGCTTACCGTTACCCAGGACACCAGCCTCCAG**
**GACGGCTGCTTGATCTACAACGTCAAGCTCAGAGGGGTGAACTTCCCATCCAACGGCCCTGTGATGCAG**
**AAGAAAACACTCGGCTGGGAGGCCACCACCGAGACCCTGTACCCCGCTGACGGCGGCCTGGAAGGCAGA**
**TGCGACATGGCCCTGAAGCTCGTGGGCGGGGGCCACCTGCACTGCAACCTGAAGACCACATACAGATCC**
**AAGAAACCCGCTAAGAACCTCAAGATGCCCGGCGTCTACTTTGTGGACCGCAGACTGGAAAGAATCAAG**
**GAGGCCGACAATGAGACCTACGTCGAGCAGCACGAGGTGGCTGTGGCCAGATACTGCGACCTCCCTAGC**
**AAACTGGGGCACAAACTTAATGGCATGGACGAGCTGTACAAGTCTGGCAGCGGAGGCTCTGGAGGCAAA**
**GAAAAGATGAGCAAAGATGGTAAAAAGAAGAAAAAGAAGTCAAAGACAAAGTGTGTAATTATGTGA**

#### Supplementary Note 7. Sequence of barcoding protein C2 (iRFP702-CAAX)

| | |
|---|---|
| **ATGGCGCGT…** | **iRFP702** |
| **TCTGGCAGC…** | **Spacer** |
| **AAAGAAAAG…** | **CAAX** |
| **TGA** | **Stop** |

**ATGGCGCGTAAGGTCGATCTCACCTCCTGCGATCGCGAGCCGATCCACATCCCCGGCAGCATTCAGCCG**
**TGCGGCTGCCTGCTAGCCTGCGACGCGCAGGCGGTGCGGATCACGCGCATTACGGAAAATGCCGGCGCG**
**TTCTTTGGACGCGAAACTCCGCGGGTCGGTGAGCTACTCGCCGATTACTTCGGCGAGACCGAAGCCCAT**
**GCGCTGCGCAACGCACTGGCGCAGTCCTCCGATCCAAAGCGACCGGCGCTGATCTTCGGTTGGCGCGAC**
**GGCCTGACCGGCCGCACCTTCGACATCTCGCTGCATCGCCATGACGGTACATCGATCATCGAGTTCGAG**
**CCTGCGGCGGCCGAACAGGCCGACAATCCGCTGCGGCTGACGCGGCAGATCATCGCGCGCACCAAAGAA**
**CTGAAGTCGCTCGAAGAGATGGCCGCACGGGTGCCGCGCTATCTGCAGGCGATGCTCGGCTATCACCGC**
**GTGATGTTGTACCGCTTCGCGGACGACGGCTCCGGCAAAGTGATCGGCGAGGCGAAGCGCAGCGACCTC**
**GAGAGCTTTCTCGGTCAGCACTTTCCGGCGTCGCTGGTCCCGCAGCAGGCGCGGCTACTGTACTTGAAG**
**AACGCGATCCGCGTGGTCTCGGATTCGCGCGGCATCAGCAGCCGGATCGTGCCCGAGCACGACGCCTCC**
**GGCGCCGCGCTTGATCTGTCGTTCGCGCACCTGCGCAGCATCTCGCCTATCCATCTCGAATTTCTGCGG**
**AACATGGGCGTCAGCGCCTCGATGTCGCTGTCGATCATCATTGACGGCACGCTATGGGGATTGATCATC**
**TGTCATCATTACGAGCCGCGTGCCGTGCCGATGGCGCAGCGCGTCGCGGCCGAAATGTTCGCCGACTTC**
**TTATCGCTGCACTTCACCGCCGCCCACCACCAACGCTCTGGCAGCGGAGGCTCTGGAGGCAAAGAAAAG**
**ATGAGCAAAGATGGTAAAAAGAAGAAAAAGAAGTCAAAGACAAAGTGTGTAATTATGTGA**

#### Supplementary Note 8. Sequence of barcoding protein D2 (BFP-CAAX)

| | |
|---|---|
| ATGGTGAGC… | BFP |
| TCTGGCAGC… | Spacer |
| AAAGAAAAG… | CAAX |
| TGA | Stop |

ATGGTGAGCAAGGGCGAGGAGCTGTTCACCGGGGTGGTGCCCATCCTGGTCGAGCTGGACGGCGACGTA
AACGGCCACAAGTTCAGCGTGAGGGGCGAGGGCGAGGGCGATGCCACCAACGGCAAGCTGACCCTGAAG
TTCATCTGCACCACCGGCAAGCTGCCCGTGCCCTGGCCCACCCTCGTGACCACCCTGAGCCACGGCGTG
CAGTGCTTCGCCCGCTACCCCGACCACATGAAGCAGCACGACTTCTTCAAGTCCGCCATGCCCGAAGGC
TACGTCCAGGAGCGCACCATCTTCTTCAAGGACGACGGCACCTACAAGACCCGCGCCGAGGTGAAGTTC
GAGGGCGACACCCTGGTGAACCGCATCGAGCTGAAGGGCGTCGACTTCAAGGAGGACGGCAACATCCTG
GGGCACAAGCTGGAGTACAACTTCAACAGCCACAACATCTATATCATGGCCGTCAAGCAGAAGAACGGC
ATCAAGGTGAACTTCAAGATCCGCCACAACGTGGAGGACGGCAGCGTGCAGCTCGCCGACCACTACCAG
CAGAACACCCCCATCGGCGACGGCCCCGTGCTGCTGCCCGACAGCCACTACCTGAGCACCCAGTCCGTG
CTGAGCAAAGACCCCAACGAGAAGCGCGATCACATGGTCCTGCTGGAGTTCCGCACCGCCGCCGGGATC
ACTCTCGGCATGGACGAGCTGTACAAGTCTGGCAGCGGAGGCTCTGGAGGCAAAGAAAAGATGAGCAAA
GATGGTAAAAAGAAGAAAAAGAAGTCAAAGACAAAGTGTGTAATTATGTGA

#### Supplementary Note 9. Sequence of barcoding protein A3 (TagRFP-LaminB1)

| | |
|---|---|
| ATGGTGTCA… | TagRFP |
| TCTGGCAGC… | Spacer |
| ATGGCGACT… | Lamin B1 |
| TGA | Stop |

ATGGTGTCAAAAGGTGAAGAACTGATTAAGGAAAATATGCACATGAAGTTATACATGGAAGGCACGGTC
AATAATCACCACTTTAAATGTACATCTGAAGGAGAGGGAAAACCGTACGAAGGAACTCAGACTATGCGA
ATCAAAGTCGTGGAGGGTGGCCCGCTGCCTTTCGCCTTCGATATTTTGGCTACTAGCTTCATGTACGGA
TCTCGCACCTTTATTAACCATACCCAGGGAATCCCAGATTTCTTTAAACAATCGTTCCCGGAAGGTTTC
ACGTGGGAAAGAGTGACGACATACGAAGATGGAGGAGTCCTCACCGCTACCCAGGACACGAGTTTGCAA
GATGGATGCTTGATCTATGATGTAAAGATTCGAGGCGTAAATTTCCCTTCGAACGGCCCAGTCATGCAA
AAGAAGACTTTGGGATGGGAAGCAAATACTGAAATGCTTTATCCTGCTGATGGAGGTCTTGAAGGACGT
ACGGATATGGCACTGAAACTGGTGGGCGGAGGACACTTGATATGTAATTTCAAAACTACGTATCGAAGT
AAAAAGCCGGCCAAGAATCTGAAAATGCCAGGAGTTTATTACGTTGACCACAGACTTGAAAGAATTAAG
GAGGCAGATAAAGAAACGTACGTGGAACAACACGAAGTAGCCGTTGCTCGCTATTGTGACTTGCCATCG
AAACTTGGACATAAACTTAATGGAATGGATGAACTTTATAAATCTGGCAGCGGAGGCTCTGGAGGCATG
GCGACTGCGACCCCCGTGCCGCCGCGGATGGGCAGCCGCGCTGGCGGCCCCACCACGCCGCTGAGCCCC
ACGCGCCTGTCGCGGCTCCAGGAGAAGGAGGAGCTGCGCGAGCTCAATGACCGGCTGGCGGTGTACATC
GACAAGGTGCGCAGCCTGGAGACGGAGAACAGCGCGCTGCAGCTGCAGGTGACGGAGCGCGAGGAGGTG
CGCGGCCGTGAGCTCACCGGCCTCAAGGCGCTCTACGAGACCGAGCTGGCCGACGCGCGACGCGCGCTC
GACGACACGGCCCGCGAGCGCGCCAAGCTGCAGATCGAGCTGGGCAAGTGCAAGGCGGAACACGACCAG
CTGCTCCTCAACTATGCTAAGAAGGAATCTGATCTTAATGGCGCCCAGATCAAGCTTCGAGAATATGAA
GCAGCACTGAATTCGAAAGATGCAGCTCTTGCTACTGCACTTGGTGACAAAAAAAGTTTAGAGGGAGAT
TTGGAGGATCTGAAGGATCAGATTGCCCAGTTGGAAGCCTCCTTAGCTGCAGCCAAAAAACAGTTAGCA
GATGAAACTTTACTTAAAGTAGATTTGGAGAATCGTTGTCAGAGCCTTACTGAGGACTTGGAGTTTCGC
AAAAGCATGTATGAAGAGGAGATTAACGAGACCAGAAGGAAGCATGAAACGCGCTTGGTAGAGGTGGAT
TCTGGGCGTCAAATTGAGTATGAGTACAAGCTGGCGCAAGCCCTTCATGAGATGAGAGAGCAACATGAT
GCCCAAGTGAGGCTGTATAAGGAGGAGCTGGAGCAGACTTACCATGCCAAACTTGAGAATGCCAGACTG
TCATCAGAGATGAATACTTCTACTGTCAACAGTGCCAGGGAAGAACTGATGGAAAGCCGCATGAGAATT
GAGAGCCTTTCATCCCAGCTTTCTAATCTACAGAAAGAGTCTAGAGCATGTTTGGAAAGGATTCAAGAA
TTAGAGGACTTGCTTGCTAAAGAAAAAGACAACTCTCGTCGCATGCTGACAGACAAAGAGAGAGAGATG
GCGGAAATAAGGGATCAAATGCAGCAACAGCTGAATGACTATGAACAGCTTCTTGATGTAAAGTTAGCC
CTGGACATGGAAATCAGTGCTTACAGGAAACTCTTAGAAGGCGAAGAAGAGAGGTTGAAGCTGTCTCCA
AGCCCTTCTTCCCGTGTGACAGTATCCCGAGCATCCTCAAGTCGTAGTGTACGTACAACTAGAGGAAAG
CGGAAGAGGGTTGATGTGGAAGAATCAGAGGCGAGTAGTAGTGTTAGCATCTCTCATTCCGCCTCAGCC
ACTGGAAATGTTTGCATCGAAGAAATTGATGTTGATGGAAATTTATCCGCTTGAAGAACACTTCTGAA
CAGGATCAACCAATGGGAGGCTGGGAGATGATCAGAAAAATTGGAGACACATCAGTCAGTTATAAATAT
ACCTCAAGATATGTGCTGAAGGCAGGCCAGACTGTTACAATTTGGGCTGCAAACGCTGGTGTCACAGCC
AGCCCCCCAACTGACCTCATCTGGAAGAACCAGAACTCGTGGGGCACTGGCGAAGATGTGAAGGTTATA
TTGAAAAATTCTCAGGGAGAGGAGGTTGCTCAAAGAAGTACAGTCTTTAAAACAACCATACCTGAAGAA
GAGGAGGAGGAGGAAGAAGCAGCTGGAGTGGTTGTTGAGGAAGAACTTTTCCACCAGCAGGGAACCCCA
AGAGCATCCAATAGAAGCTGTGCAATTATGTGA

#### Supplementary Note 10. Sequence of barcoding protein B3 (mCardinal-LaminB1)

| | |
|---|---|
| ATGGTGAGC… | mCardinal |
| TCTGGCAGC… | Spacer |
| ATGGCGACT… | Lamin B1 |
| TGA | Stop |

ATGGTGAGCAAGGGCGAGGAGCTGATCAAGGAGAACATGCACATGAAGCTGTACATGGAAGGCACCGTG
AACAACCACCACTTCAAGTGCACCACCGAAGGGGAGGGCAAGCCCTACGAGGGCACCCAGACCCAGAGG
ATTAAGGTGGTGGAGGGAGGCCCCCTGCCGTTCGCATTCGACATCCTGGCCACCTGCTTTATGTACGGG
AGCAAGACCTTCATCAACCACACCCAGGGCATCCCCGATTTCTTTAAGCAGTCCTTCCCTGAGGGCTTC
ACATGGGAGAGAGTCACCACATACGAAGACGGGGGCGTGCTTACCGTTACCCAGGACACCAGCCTCCAG
GACGGCTGCTTGATCTACAACGTCAAGCTCAGAGGGGTGAACTTCCCATCCAACGGCCCTGTGATGCAG
AAGAAAACACTCGGCTGGGAGGCCACCACCGAGACCCTGTACCCCGCTGACGGCGGCCTGGAAGGCAGA
TGCGACATGGCCCTGAAGCTCGTGGGCGGGGGCCACCTGCACTGCAACCTGAAGACCACATACAGATCC
AAGAAACCCGCTAAGAACCTCAAGATGCCCGGCGTCTACTTTGTGGACCGCAGACTGGAAAGAATCAAG
GAGGCCGACAATGAGACCTACGTCGAGCAGCACGAGGTGGCTGTGGCCAGATACTGCGACCTCCCTAGC
AAACTGGGGCACAAACTTAATGGCATGGACGAGCTGTACAAGTCTGGCAGCGGAGGCTCTGGAGGCATG
GCGACTGCGACCCCCGTGCCGCCGCGGATGGGCAGCCGCGCTGGCGGCCCCACCACGCCGCTGAGCCCC
ACGCGCCTGTCGCGGCTCCAGGAGAAGGAGGAGCTGCGCGAGCTCAATGACCGGCTGGCGGTGTACATC
GACAAGGTGCGCAGCCTGGAGACGGAGAACAGCGCGCTGCAGCTGCAGGTGACGGAGCGCGAGGAGGTG
CGCGGCCGTGAGCTCACCGGCCTCAAGGCGCTCTACGAGACCGAGCTGGCCGACGCGCGACGCGCGCTC
GACGACACGGCCCGCGAGCGCGCCAAGCTGCAGATCGAGCTGGGCAAGTGCAAGGCGGAACACGACCAG
CTGCTCCTCAACTATGCTAAGAAGGAATCTGATCTTAATGGCGCCCAGATCAAGCTTCGAGAATATGAA
GCAGCACTGAATTCGAAAGATGCAGCTCTTGCTACTGCACTTGGTGACAAAAAAAGTTTAGAGGGAGAT
TTGGAGGATCTGAAGGATCAGATTGCCCAGTTGGAAGCCTCCTTAGCTGCAGCCAAAAAACAGTTAGCA
GATGAAACTTTACTTAAAGTAGATTTGGAGAATCGTTGTCAGAGCCTTACTGAGGACTTGGAGTTTCGC
AAAAGCATGTATGAAGAGGAGATTAACGAGACCAGAAGGAAGCATGAAACGCGCTTGGTAGAGGTGGAT
TCTGGGCGTCAAATTGAGTATGAGTACAAGCTGGCGCAAGCCCTTCATGAGATGAGAGAGCAACATGAT
GCCCAAGTGAGGCTGTATAAGGAGGAGCTGGAGCAGACTTACCATGCCAAACTTGAGAATGCCAGACTG
TCATCAGAGATGAATACTTCTACTGTCAACAGTGCCAGGGAAGAACTGATGGAAAGCCGCATGAGAATT
GAGAGCCTTTCATCCCAGCTTTCTAATCTACAGAAAGAGTCTAGAGCATGTTTGGAAAGGATTCAAGAA
TTAGAGGACTTGCTTGCTAAAGAAAAAGACAACTCTCGTCGCATGCTGACAGACAAAGAGAGAGAGATG
GCGGAAATAAGGGATCAAATGCAGCAACAGCTGAATGACTATGAACAGCTTCTTGATGTAAAGTTAGCC
CTGGACATGGAAATCAGTGCTTACAGGAAACTCTTAGAAGGCGAAGAAGAGAGGTTGAAGCTGTCTCCA
AGCCCTTCTTCCCGTGTGACAGTATCCCGAGCATCCTCAAGTCGTAGTGTACGTACAACTAGAGGAAAG
CGGAAGAGGGTTGATGTGGAAGAATCAGAGGCGAGTAGTAGTGTTAGCATCTCTCATTCCGCCTCAGCC
ACTGGAAATGTTTGCATCGAAGAAATTGATGTTGATGGAAATTTATCCGCTTGAAGAACACTTCTGAA
CAGGATCAACCAATGGGAGGCTGGGAGATGATCAGAAAAATTGGAGACACATCAGTCAGTTATAAATAT
ACCTCAAGATATGTGCTGAAGGCAGGCCAGACTGTTACAATTTGGGCTGCAAACGCTGGTGTCACAGCC
AGCCCCCCAACTGACCTCATCTGGAAGAACCAGAACTCGTGGGGCACTGGCGAAGATGTGAAGGTTATA
TTGAAAAATTCTCAGGGAGAGGAGGTTGCTCAAAGAAGTACAGTCTTTAAAACAACCATACCTGAAGAA
GAGGAGGAGGAGGAAGAAGCAGCTGGAGTGGTTGTTGAGGAAGAACTTTTCCACCAGCAGGGAACCCCA
AGAGCATCCAATAGAAGCTGTGCAATTATGTGA

#### Supplementary Note 11. Sequence of barcoding protein C3 (iRFP702-LaminB1)

| | |
|---|---|
| ATGGCGCGT… | iRFP702 |
| TCTGGCAGC… | Spacer |
| ATGGCGACT… | Lamin B1 |
| TGA | Stop |

ATGGCGCGTAAGGTCGATCTCACCTCCTGCGATCGCGAGCCGATCCACATCCCCGGCAGCATTCAGCCG
TGCGGCTGCCTGCTAGCCTGCGACGCGCAGGCGGTGCGGATCACGCGCATTACGGAAAATGCCGGCGCG
TTCTTTGGACGCGAAACTCCGCGGGTCGGTGAGCTACTCGCCGATTACTTCGGCGAGACCGAAGCCCAT
GCGCTGCGCAACGCACTGGCGCAGTCCTCCGATCCAAAGCGACCGGCGCTGATCTTCGGTTGGCGCGAC
GGCCTGACCGGCCGCACCTTCGACATCTCGCTGCATCGCCATGACGGTACATCGATCATCGAGTTCGAG
CCTGCGGCGGCCGAACAGGCCGACAATCCGCTGCGGCTGACGCGGCAGATCATCGCGCGCACCAAAGAA
CTGAAGTCGCTCGAAGAGATGGCCGCACGGGTGCCGCGCTATCTGCAGGCGATGCTCGGCTATCACCGC
GTGATGTTGTACCGCTTCGCGGACGACGGCTCCGGCAAAGTGATCGGCGAGGCGAAGCGCAGCGACCTC
GAGAGCTTTCTCGGTCAGCACTTTCCGGCGTCGCTGGTCCCGCAGCAGGCGCGGCTACTGTACTTGAAG
AACGCGATCCGCGTGGTCTCGGATTCGCGCGGCATCAGCAGCCGGATCGTGCCCGAGCACGACGCCTCC
GGCGCCGCGCTTGATCTGTCGTTCGCGCACCTGCGCAGCATCTCGCCTATCCATCTCGAATTTCTGCGG
AACATGGGCGTCAGCGCCTCGATGTCGCTGTCGATCATCATTGACGGCACGCTATGGGGATTGATCATC
TGTCATCATTACGAGCCGCGTGCCGTGCCGATGGCGCAGCGCGTCGCGGCCGAAATGTTCGCCGACTTC
TTATCGCTGCACTTCACCGCCGCCCACCACCAACGCTCTGGCAGCGGAGGCTCTGGAGGCATGGCGACT
GCGACCCCCGTGCCGCCGCGGATGGGCAGCCGCGCTGGCGGCCCCACCACGCCGCTGAGCCCCACGCGC
CTGTCGCGGCTCCAGGAGAAGGAGGAGCTGCGCGAGCTCAATGACCGGCTGGCGGTGTACATCGACAAG
GTGCGCAGCCTGGAGACGGAGAACAGCGCGCTGCAGCTGCAGGTGACGGAGCGCGAGGAGGTGCGCGGC
CGTGAGCTCACCGGCCTCAAGGCGCTCTACGAGACCGAGCTGGCCGACGCGCGACGCGCGCTCGACGAC
ACGGCCCGCGAGCGCGCCAAGCTGCAGATCGAGCTGGGCAAGTGCAAGGCGGAACACGACCAGCTGCTC
CTCAACTATGCTAAGAAGGAATCTGATCTTAATGGCGCCCAGATCAAGCTTCGAGAATATGAAGCAGCA
CTGAATTCGAAAGATGCAGCTCTTGCTACTGCACTTGGTGACAAAAAAAGTTTAGAGGGAGATTTGGAG
GATCTGAAGGATCAGATTGCCCAGTTGGAAGCCTCCTTAGCTGCAGCCAAAAAACAGTTAGCAGATGAA
ACTTTACTTAAAGTAGATTTGGAGAATCGTTGTCAGAGCCTTACTGAGGACTTGGAGTTTCGCAAAAGC
ATGTATGAAGAGGAGATTAACGAGACCAGAAGGAAGCATGAAACGCGCTTGGTAGAGGTGGATTCTGGG
CGTCAAATTGAGTATGAGTACAAGCTGGCGCAAGCCCTTCATGAGATGAGAGAGCAACATGATGCCCAA
GTGAGGCTGTATAAGGAGGAGCTGGAGCAGACTTACCATGCCAAACTTGAGAATGCCAGACTGTCATCA
GAGATGAATACTTCTACTGTCAACAGTGCCAGGGAAGAACTGATGGAAAGCCGCATGAGAATTGAGAGC
CTTTCATCCCAGCTTTCTAATCTACAGAAAGAGTCTAGAGCATGTTTGGAAAGGATTCAAGAATTAGAG
GACTTGCTTGCTAAAGAAAAAGACAACTCTCGTCGCATGCTGACAGACAAAGAGAGAGAGATGGCGGAA
ATAAGGGATCAAATGCAGCAACAGCTGAATGACTATGAACAGCTTCTTGATGTAAAGTTAGCCCTGGAC
ATGGAAATCAGTGCTTACAGGAAACTCTTAGAAGGCGAAGAAGAGAGGTTGAAGCTGTCTCCAAGCCCT
TCTTCCCGTGTGACAGTATCCCGAGCATCCTCAAGTCGTAGTGTACGTACAACTAGAGGAAAGCGGAAG
AGGGTTGATGTGGAAGAATCAGAGGCGAGTAGTAGTGTTAGCATCTCTCATTCCGCCTCAGCCACTGGA
AATGTTTGCATCGAAGAAATTGATGTTGATGGAAATTTATCCGCTTGAAGAACACTTCTGAACAGGAT
CAACCAATGGGAGGCTGGGAGATGATCAGAAAAATTGGAGACACATCAGTCAGTTATAAATATACCTCA
AGATATGTGCTGAAGGCAGGCCAGACTGTTACAATTTGGGCTGCAAACGCTGGTGTCACAGCCAGCCCC
CCAACTGACCTCATCTGGAAGAACCAGAACTCGTGGGGCACTGGCGAAGATGTGAAGGTTATATTGAAA
AATTCTCAGGGAGAGGAGGTTGCTCAAAGAAGTACAGTCTTTAAAACAACCATACCTGAAGAAGAGGAG
GAGGAGGAAGAAGCAGCTGGAGTGGTTGTTGAGGAAGAACTTTTCCACCAGCAGGGAACCCCAAGAGCA
TCCAATAGAAGCTGTGCAATTATGTGA

#### Supplementary Note 12. Sequence of barcoding protein D3 (BFP-LaminB1)

| | |
|---|---|
| ATGGTGAGC… | BFP |
| TCTGGCAGC… | Spacer |
| ATGGCGACT… | Lamin B1 |
| TGA | Stop |

ATGGTGAGCAAGGGCGAGGAGCTGTTCACCGGGGTGGTGCCCATCCTGGTCGAGCTGGACGGCGACGTA
AACGGCCACAAGTTCAGCGTGAGGGGCGAGGGCGAGGGCGATGCCACCAACGGCAAGCTGACCCTGAAG
TTCATCTGCACCACCGGCAAGCTGCCCGTGCCCTGGCCCACCCTCGTGACCACCCTGAGCCACGGCGTG
CAGTGCTTCGCCCGCTACCCCGACCACATGAAGCAGCACGACTTCTTCAAGTCCGCCATGCCCGAAGGC
TACGTCCAGGAGCGCACCATCTTCTTCAAGGACGACGGCACCTACAAGACCCGCGCCGAGGTGAAGTTC
GAGGGCGACACCCTGGTGAACCGCATCGAGCTGAAGGGCGTCGACTTCAAGGAGGACGGCAACATCCTG
GGGCACAAGCTGGAGTACAACTTCAACAGCCACAACATCTATATCATGGCCGTCAAGCAGAAGAACGGC
ATCAAGGTGAACTTCAAGATCCGCCACAACGTGGAGGACGGCAGCGTGCAGCTCGCCGACCACTACCAG
CAGAACACCCCCATCGGCGACGGCCCCGTGCTGCTGCCCGACAGCCACTACCTGAGCACCCAGTCCGTG
CTGAGCAAAGACCCCAACGAGAAGCGCGATCACATGGTCCTGCTGGAGTTCCGCACCGCCGCCGGGATC
ACTCTCGGCATGGACGAGCTGTACAAGTCTGGCAGCGGAGGCTCTGGAGGCATGGCGACTGCGACCCCC
GTGCCGCCGCGGATGGGCAGCCGCGCTGGCGGCCCCACCACGCCGCTGAGCCCCACGCGCCTGTCGCGG
CTCCAGGAGAAGGAGGAGCTGCGCGAGCTCAATGACCGGCTGGCGGTGTACATCGACAAGGTGCGCAGC
CTGGAGACGGAGAACAGCGCGCTGCAGCTGCAGGTGACGGAGCGCGAGGAGGTGCGCGGCCGTGAGCTC
ACCGGCCTCAAGGCGCTCTACGAGACCGAGCTGGCCGACGCGCGACGCGCGCTCGACGACACGGCCCGC
GAGCGCGCCAAGCTGCAGATCGAGCTGGGCAAGTGCAAGGCGGAACACGACCAGCTGCTCCTCAACTAT
GCTAAGAAGGAATCTGATCTTAATGGCGCCCAGATCAAGCTTCGAGAATATGAAGCAGCACTGAATTCG
AAAGATGCAGCTCTTGCTACTGCACTTGGTGACAAAAAAAGTTTAGAGGGAGATTTGGAGGATCTGAAG
GATCAGATTGCCCAGTTGGAAGCCTCCTTAGCTGCAGCCAAAAAACAGTTAGCAGATGAAACTTTACTT
AAAGTAGATTTGGAGAATCGTTGTCAGAGCCTTACTGAGGACTTGGAGTTTCGCAAAAGCATGTATGAA
GAGGAGATTAACGAGACCAGAAGGAAGCATGAAACGCGCTTGGTAGAGGTGGATTCTGGGCGTCAAATT
GAGTATGAGTACAAGCTGGCGCAAGCCCTTCATGAGATGAGAGAGCAACATGATGCCCAAGTGAGGCTG
TATAAGGAGGAGCTGGAGCAGACTTACCATGCCAAACTTGAGAATGCCAGACTGTCATCAGAGATGAAT
ACTTCTACTGTCAACAGTGCCAGGGAAGAACTGATGGAAAGCCGCATGAGAATTGAGAGCCTTTCATCC
CAGCTTTCTAATCTACAGAAAGAGTCTAGAGCATGTTTGGAAAGGATTCAAGAATTAGAGGACTTGCTT
GCTAAAGAAAAAGACAACTCTCGTCGCATGCTGACAGACAAAGAGAGAGAGATGGCGGAAATAAGGGAT
CAAATGCAGCAACAGCTGAATGACTATGAACAGCTTCTTGATGTAAAGTTAGCCCTGGACATGGAAATC
AGTGCTTACAGGAAACTCTTAGAAGGCGAAGAAGAGAGGTTGAAGCTGTCTCCAAGCCCTTCTTCCCGT
GTGACAGTATCCCGAGCATCCTCAAGTCGTAGTGTACGTACAACTAGAGGAAAGCGGAAGAGGGTTGAT
GTGGAAGAATCAGAGGCGAGTAGTAGTGTTAGCATCTCTCATTCCGCCTCAGCCACTGGAAATGTTTGC
ATCGAAGAAATTGATGTTGATGGAAATTTATCCGCTTGAAGAACACTTCTGAACAGGATCAACCAATG
GGAGGCTGGGAGATGATCAGAAAAATTGGAGACACATCAGTCAGTTATAAATATACCTCAAGATATGTG
CTGAAGGCAGGCCAGACTGTTACAATTTGGGCTGCAAACGCTGGTGTCACAGCCAGCCCCCCAACTGAC
CTCATCTGGAAGAACCAGAACTCGTGGGGCACTGGCGAAGATGTGAAGGTTATATTGAAAAATTCTCAG
GGAGAGGAGGTTGCTCAAAGAAGTACAGTCTTTAAAACAACCATACCTGAAGAAGAGGAGGAGGAGGAA
GAAGCAGCTGGAGTGGTTGTTGAGGAAGAACTTTTCCACCAGCAGGGAACCCCAAGAGCATCCAATAGA
AGCTGTGCAATTATGTGA

#### Supplementary Note 13. Sequence of barcoding protein A4 (TagRFP-NES)

| | |
|---|---|
| ATGGTGTCA… | TagRFP |
| TCTGGCAGC… | Spacer |
| CTCCAGAAG… | MAPKK NES |
| TGA | Stop |

ATGGTGTCAAAAGGTGAAGAACTGATTAAGGAAAATATGCACATGAAGTTATACATGGAAGGCACGGTC
AATAATCACCACTTTAAATGTACATCTGAAGGAGAGGGAAAACCGTACGAAGGAACTCAGACTATGCGA
ATCAAAGTCGTGGAGGGTGGCCCGCTGCCTTTCGCCTTCGATATTTTGGCTACTAGCTTCATGTACGGA
TCTCGCACCTTTATTAACCATACCCAGGGAATCCCAGATTTCTTTAAACAATCGTTCCCGGAAGGTTTC
ACGTGGGAAAGAGTGACGACATACGAAGATGGAGGAGTCCTCACCGCTACCCAGGACACGAGTTTGCAA
GATGGATGCTTGATCTATGATGTAAAGATTCGAGGCGTAAATTTCCCTTCGAACGGCCCAGTCATGCAA
AAGAAGACTTTGGGATGGGAAGCAAATACTGAAATGCTTTATCCTGCTGATGGAGGTCTTGAAGGACGT
ACGGATATGGCACTGAAACTGGTGGGCGGAGGACACTTGATATGTAATTTCAAAACTACGTATCGAAGT
AAAAAGCCGGCCAAGAATCTGAAAATGCCAGGAGTTTATTACGTTGACCACAGACTTGAAAGAATTAAG
GAGGCAGATAAAGAAACGTACGTGGAACAACACGAAGTAGCCGTTGCTCGCTATTGTGACTTGCCATCG
AAACTTGGACATAAACTTAATGGAATGGATGAACTTTATAAAggaagctccCTCCAGAAGAAACTGGAG
GAACTGGAACTGTGA

#### Supplementary Note 14. Sequence of barcoding protein B4 (mCardinal-NES)

| | |
|---|---|
| **ATGGTGAGC…** | **mCardinal** |
| **TCTGGCAGC…** | **Spacer** |
| **CTCCAGAAG…** | **MAPKK NES** |
| **TGA** | **Stop** |

**ATGGTGAGCAAGGGCGAGGAGCTGATCAAGGAGAACATGCACATGAAGCTGTACATGGAAGGCACCGTG
AACAACCACCACTTCAAGTGCACCACCGAAGGGGAGGGCAAGCCCTACGAGGGCACCCAGACCCAGAGG
ATTAAGGTGGTGGAGGGAGGCCCCCTGCCGTTCGCATTCGACATCCTGGCCACCTGCTTTATGTACGGG
AGCAAGACCTTCATCAACCACACCCAGGGCATCCCCGATTTCTTTAAGCAGTCCTTCCCTGAGGGCTTC
ACATGGGAGAGAGTCACCACATACGAAGACGGGGGCGTGCTTACCGTTACCCAGGACACCAGCCTCCAG
GACGGCTGCTTGATCTACAACGTCAAGCTCAGAGGGGTGAACTTCCCATCCAACGGCCCTGTGATGCAG
AAGAAAACACTCGGCTGGGAGGCCACCACCGAGACCCTGTACCCCGCTGACGGCGGCCTGGAAGGCAGA
TGCGACATGGCCCTGAAGCTCGTGGGCGGGGGCCACCTGCACTGCAACCTGAAGACCACATACAGATCC
AAGAAACCCGCTAAGAACCTCAAGATGCCCGGCGTCTACTTTGTGGACCGCAGACTGGAAAGAATCAAG
GAGGCCGACAATGAGACCTACGTCGAGCAGCACGAGGTGGCTGTGGCCAGATACTGCGACCTCCCTAGC
AAACTGGGGCACAAACTTAATGGCATGGACGAGCTGTACAAAggaagctccCTCCAGAAGAAACTGGAG
GAACTGGAACTGTGA**

#### Supplementary Note 15. Sequence of barcoding protein C4 (iRFP702-NES)

| | |
|---|---|
| ATGGCGCGT… | iRFP702 |
| TCTGGCAGC… | Spacer |
| CTCCAGAAG… | MAPKK NES |
| TGA | Stop |

ATGGCGCGTAAGGTCGATCTCACCTCCTGCGATCGCGAGCCGATCCACATCCCCGGCAGCATTCAGCCG
TGCGGCTGCCTGCTAGCCTGCGACGCGCAGGCGGTGCGGATCACGCGCATTACGGAAAATGCCGGCGCG
TTCTTTGGACGCGAAACTCCGCGGGTCGGTGAGCTACTCGCCGATTACTTCGGCGAGACCGAAGCCCAT
GCGCTGCGCAACGCACTGGCGCAGTCCTCCGATCCAAAGCGACCGGCGCTGATCTTCGGTTGGCGCGAC
GGCCTGACCGGCCGCACCTTCGACATCTCGCTGCATCGCCATGACGGTACATCGATCATCGAGTTCGAG
CCTGCGGCGGCCGAACAGGCCGACAATCCGCTGCGGCTGACGCGGCAGATCATCGCGCGCACCAAAGAA
CTGAAGTCGCTCGAAGAGATGGCCGCACGGGTGCCGCGCTATCTGCAGGCGATGCTCGGCTATCACCGC
GTGATGTTGTACCGCTTCGCGGACGACGGCTCCGGCAAAGTGATCGGCGAGGCGAAGCGCAGCGACCTC
GAGAGCTTTCTCGGTCAGCACTTTCCGGCGTCGCTGGTCCCGCAGCAGGCGCGGCTACTGTACTTGAAG
AACGCGATCCGCGTGGTCTCGGATTCGCGCGGCATCAGCAGCCGGATCGTGCCCGAGCACGACGCCTCC
GGCGCCGCGCTTGATCTGTCGTTCGCGCACCTGCGCAGCATCTCGCCTATCCATCTCGAATTTCTGCGG
AACATGGGCGTCAGCGCCTCGATGTCGCTGTCGATCATCATTGACGGCACGCTATGGGGATTGATCATC
TGTCATCATTACGAGCCGCGTGCCGTGCCGATGGCGCAGCGCGTCGCGGCCGAAATGTTCGCCGACTTC
TTATCGCTGCACTTCACCGCCGCCCACCACCAACGCggaagctccCTCCAGAAGAAACTGGAGGAACTG
GAACTGTGA

#### Supplementary Note 16. Sequence of barcoding protein D4 (BFP-NES)

| | |
|---|---|
| **ATGGTGAGC…** | **BFP** |
| **TCTGGCAGC…** | **Spacer** |
| **CTCCAGAAG…** | **MAPKK NES** |
| **TGA** | **Stop** |

**ATGGTGAGCAAGGGCGAGGAGCTGTTCACCGGGGTGGTGCCCATCCTGGTCGAGCTGGACGGCGACGTA**
**AACGGCCACAAGTTCAGCGTGAGGGGCGAGGGCGAGGGCGATGCCACCAACGGCAAGCTGACCCTGAAG**
**TTCATCTGCACCACCGGCAAGCTGCCCGTGCCCTGGCCCACCCTCGTGACCACCCTGAGCCACGGCGTG**
**CAGTGCTTCGCCCGCTACCCCGACCACATGAAGCAGCACGACTTCTTCAAGTCCGCCATGCCCGAAGGC**
**TACGTCCAGGAGCGCACCATCTTCTTCAAGGACGACGGCACCTACAAGACCCGCGCCGAGGTGAAGTTC**
**GAGGGCGACACCCTGGTGAACCGCATCGAGCTGAAGGGCGTCGACTTCAAGGAGGACGGCAACATCCTG**
**GGGCACAAGCTGGAGTACAACTTCAACAGCCACAACATCTATATCATGGCCGTCAAGCAGAAGAACGGC**
**ATCAAGGTGAACTTCAAGATCCGCCACAACGTGGAGGACGGCAGCGTGCAGCTCGCCGACCACTACCAG**
**CAGAACACCCCCATCGGCGACGGCCCCGTGCTGCTGCCCGACAGCCACTACCTGAGCACCCAGTCCGTG**
**CTGAGCAAAGACCCCAACGAGAAGCGCGATCACATGGTCCTGCTGGAGTTCCGCACCGCCGCCGGGATC**
**ACTCTCGGCATGGACGAGCTGTACAAAggaagctccCTCCAGAAGAAACTGGAGGAACTGGAACTGTGA**
